# CytoCensus: mapping cell identity and division in tissues and organs using machine learning

**DOI:** 10.1101/137406

**Authors:** Martin Hailstone, Dominic Waithe, Tamsin J Samuels, Lu Yang, Ita Costello, Yoav Arava, Elizabeth J Robertson, Richard M Parton, Ilan Davis

## Abstract

A major challenge in cell and developmental biology is the automated identification and quantitation of cells in complex multilayered tissues. We developed CytoCensus: an easily deployed implementation of supervised machine learning that extends convenient 2D “point-and-click” user training to 3D detection of cells in challenging datasets with ill-defined cell boundaries. In tests on these datasets, CytoCensus outperforms other freely available image analysis software in accuracy and speed of cell detection. We used CytoCensus to count stem cells and their progeny, and to quantify individual cell divisions from time-lapse movies of explanted *Drosophila* larval brains, comparing wild-type and mutant phenotypes. We further illustrate the general utility and future potential of CytoCensus by analysing the 3D organisation of multiple cell classes in Zebrafish retinal organoids and cell distributions in mouse embryos. CytoCensus opens the possibility of straightforward and robust automated analysis of developmental phenotypes in complex tissues.

**Summary:** Hailstone *et al*. develop CytoCensus, a “point-and-click” supervised machine-learning image analysis software to quantitatively identify defined cell classes and divisions from large multidimensional data sets of complex tissues. They demonstrate its utility in analysing challenging developmental phenotypes in living explanted *Drosophila* larval brains, mammalian embryos and zebrafish organoids. They further show, in comparative tests, a significant improvement in performance over existing easy-to-use image analysis software.

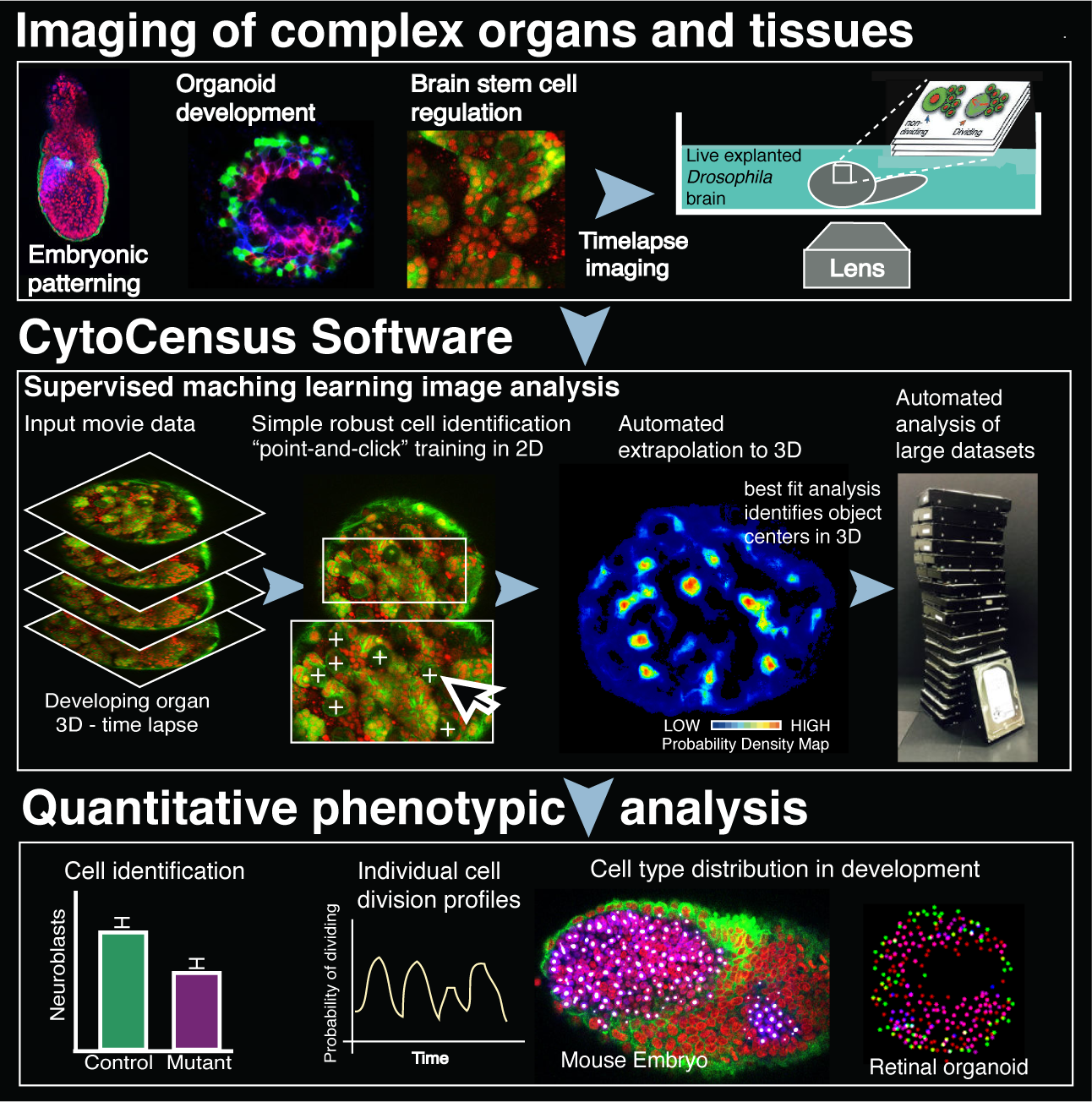

**Highlights:**

- CytoCensus: machine learning quantitation of cell types in complex 3D tissues
- Single cell analysis of division rates from movies of living *Drosophila* brains in 3D
- Diverse applications in the analysis of developing vertebrate tissues and organoids
- Outperforms other image analysis software on challenging, low SNR datasets tested

## Introduction

Complex tissues develop through regulated proliferation and differentiation of a small number of stem cells. For example, in the brain these processes of proliferation and differentiation lead to a vast and diverse population of neurons and glia from a limited number of neural stem cells, also known as neuroblasts (NBs) in *Drosophila* (Kohwi & Doe, 2013). Elucidating the molecular basis of such developmental processes is not only essential for understanding basic neuroscience, but is also important for discovering new treatments for neurological diseases and cancer. Modern imaging approaches have proven indispensable in studying development in intact zebrafish (*Danio rario*) and *Drosophila* tissues (Barbosa & Ninkovic, 2016; Dray *et al*., 2015; Medioni *et al*., 2015; Rabinovich *et al*., 2015; Cabernard & Doe., 2013; Graeden & Sive, 2009). Tissue imaging approaches have also been combined with functional genetic screens, for example to discover neuroblast (NB) behaviour underlying defects in brain size or tumour formation (Berger *et al*., 2012; Homem & Knoblich, 2012; Neumüller *et al*., 2011). Such screens have the power of genome wide coverage, but to be effective, require detailed characterisation of phenotypes using image analysis. Often these kinds of screens are limited in their power by the fact that phenotypic analysis of complex tissues can only be carried out using manual image analysis methods or complex bespoke image analysis.

*Drosophila* larval brains develop for more than 120 hours (Homem & Knoblich, 2012), a process best characterised by long term time-lapse microscopy. However, to date, imaging intact developing live brains has tended to be carried out for relatively short periods of a few hours (Lerit *et al*., 2014; Cabernard & Doe, 2013; Prithviraj *et al*., 2012) or using disaggregated brain cells in culture (Homem *et al*., 2013; Moraru *et al*., 2012; Savoian & Rieder, 2002; Furst & Mahowald, 1985). Furthermore, although extensively studied, a range of different division rates for both NB and GMCs are reported in the literature (Homem *et al*., 2013; Bowman *et al*., 2008; Ceron *et al*., 2006) and in general, division rates have not been systematically determined for individual neuroblasts. Imaging approaches have improved rapidly in speed and sensitivity, making imaging of live intact tissues in 3D possible over developmentally relevant time-scales. However, long term exposure to light often perturbs the behaviour of cells in subtle ways. Moreover, automated methods for the analysis of the resultant huge datasets are still lagging behind the microscopy methods. These imaging and analysis problems limit our ability to study NB development in larval brains, as well as more generally our ability to study complex tissues and organs.

Here, we describe our development and validation of *ex vivo* live imaging of *Drosophila* brains, and of CytoCensus, a machine learning-based automated image analysis software that fills the technology gap that exists for images of complex tissues and organs where segmentation and spot detection approaches can struggle. Our program efficiently and accurately identifies cell types and divisions of interest in very large (50 GB) multichannel 3D and 4D datasets, outperforming other state-of-the-art tools that we tested. We demonstrate the effectiveness and flexibility of CytoCensus first by quantitating cell type and division rates in *ex vivo* cultured intact developing *Drosophila* larval brains imaged at 10% of the normal illumination intensity with image quality restoration using patched-based denoising algorithms. Second, we quantitatively characterise the precise numbers and distributions of the different cell classes within two vertebrate tissues: 3D Zebrafish organoids and mouse embryos. In all these cases, CytoCensus successfully outputs quantitation of the distributions of most cells in tissues that are too large or complex for practical manual annotation. Our software provides a convenient tool that works “out-of-the-box” for quantitation and single cell analysis of complex tissues in 4D, and, in combination with other software (eg. FIJI), supports the study of more complex problems than would otherwise be possible. CytoCensus offers a practical alternative to producing bespoke image analysis pipelines for specific applications.

### Motivation and design

We sought to overcome the image analysis bottleneck that exists for complex tissues and organs by creating easy to use, automated image analysis tools able to accurately identify cell types and determine their distributions and division rates in 3D, over time within intact tissues. To date, challenging image analysis tasks of this sort have largely depended on slow, painstaking manual analysis, or the bespoke development or modification of dedicated specialised tools by an image analyst with significant programming skills (Chittajallu *et al*., 2015; Schmitz *et al*., 2014; Stegmaier *et al*., 2014; Homem *et al*., 2013; Myers 2012; Meijering, 2012; Meijering *et al*., 2012; Rittscher, 2010). Of the current freely-available automated tools, amongst the most powerful are Ilastik and the customised pipelines of the FARSIGHT toolbox and CellProfiler (Padmanabhan *et al*., 2014; Sommer,& Gerlich, 2013; Sommer *et al*., 2011; Roysam *et al*., 2008). However, these three approaches require advanced knowledge of image processing, programming and/or extensive manual annotation. Other software such as Advanced Cell Classifier are targeted at analysis of 2D data, whilst programs such as RACE, SuRVoS, 3D-RSD and MINS are generally tailored to specific applications (Luengo *et al*., 2017; Stegmaier *et al*., 2016; Lou *et al*., 2014; Cabernard & Doe, 2013; Homem *et al*., 2013; Arganda-Carreras *et al*., 2017; Logan *et al*., 2016; Gertych *et al*., 2015). Recently, efforts to make deep learning approaches easily accessible have made great strides (Falk *et al*., 2019); such implementations have the potential to increase access to these powerful supervised segmentation methods, but at present hardware and installation requirements are likely to be too complex for the typical biologist. In general, we find that existing tools can be powerful in specific examples, but lack the flexibility, speed and/or ease of use to make them effective solutions for most biologists in the analysis of large time-lapse movies of 3D developing tissues. In developing CytoCensus, we sought to design a widely applicable, supervised machine leaning-based, image analysis tool, addressing the needs of biologists to efficiently characterise and quantitate dense complex 3D tissues at the single cell level with practical imaging conditions. This level of analysis of developing tissues, organoids or organs is frequently difficult due to the complexity and density of the tissue arrangement or labelling, as well as limitations of signal to noise. We therefore aimed to make CytoCensus robust to these issues but also to make it as user friendly as possible. In contrast to other image analysis approaches that require the user to define the cell boundaries, CytoCensus simply requires the user to point-and-click on the approximate centres of cells. This single click training need only be carried out on a few representative 2D planes from a large 3D volume, and tolerates relatively poor image quality compatible with extended live cell imaging. To make the task very user friendly, we preconfigured most algorithm settings leaving a few, largely intuitive parameters, for the user to set. To improve performance, we enabled users to define regions of interest (ROIs), which exclude parts of a tissue that are not of interest or interfere with the analysis. We also separated the training phase from the analysis phase, allowing efficient batch processing of data. CytoCensus then determines the probability of each pixel in the image being the centre of the highlighted cell class in 3D, based on the characteristics of the pixels around the site clicked. This proximity map is used to identify all of the cells of interest. Finally, to increase the ease of adoption, we designed CytoCensus to be easily installed and work on multiple platforms and computers with standard specifications, including generically configured laptops without any pre-requisites. Collectively, these improvements make CytoCensus an accessible and user-friendly image analysis tool that will enable biologists to analyse their image data effectively, increase experimental throughput and increase the statistical strength of their conclusions.

## Results

### Optimised time-lapse imaging of developing intact *ex-vivo* brains

To extend our ability to study stem cell behaviour in the context of the intact *Drosophila* brain, we modified the methods of Cabernard & Doe (2013), revised in Syed *et al*., (2017), to produce a convenient and effective protocol optimising tissue viability for long-term culture and quantitative imaging. We first developed an isolation procedure incorporating scissor-based dissection of second or third-instar larvae, in preference to solely tweezer or needle-based dissection which can damage the tissue. We then simplified the culture medium and developed a convenient brain mounting technique that immobilises the organ using agar (Figure 1A; Materials & Methods). We also made use of bright, endogenously expressed fluorescently tagged proteins Jupiter::GFP and Histone::RFP marking microtubules and chromosomes respectively, to follow the developing brain (Figure 1B). We chose generic cytological markers as these are more consistent across *wild-type* (WT) and different mutants than more specific markers, such as Deadpan (Dpn), Asense (Ase) or Prospero (Pros), commonly used to identify NBs, GMCs and neurons. Finally, we optimised the imaging conditions to provide 3D data sets of sufficient temporal and spatial resolution to follow cell proliferation over time without compromising viability (see Materials & Methods). Significantly, to maximise temporal and spatial resolution without causing damage, we reduced photo-damage by decreasing the laser excitation power by approximately 10 fold (see Materials & Methods) and subsequently restoring image quality using patch-based denoising (Carlton *et al*., 2010), developed by Kervrann and Boulanger (2006). This approach allowed us to follow the lineage and quantitate the divisions of NB and GMCs in the intact brain in 3D (Figure 1C, D).

**Figure 1.**
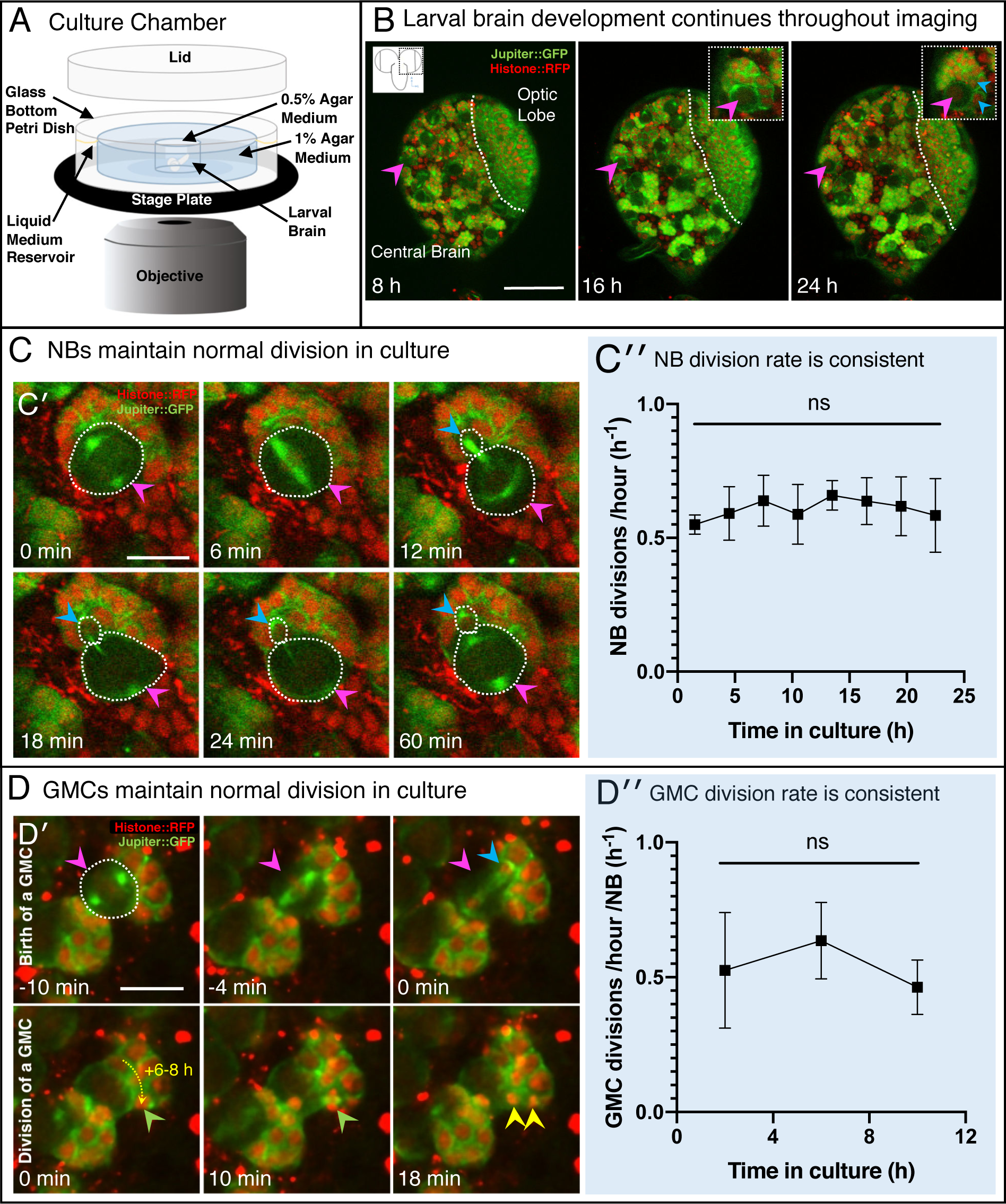
Extended 3D time-lapse imaging of live *ex-vivo* cultured brains. **A**) Diagram of the chamber and sample preparation for long-term time-lapse imaging on an inverted microscope (see Materials & Methods). **B**) 24 h, confocal 3D time-lapse imaging of a developing larval brain lobe (inset, top left, shows orientation and region of the brain imaged) labelled with Jupiter::GFP and Histone::RFP, and registered over time to account for movement. Arrowheads indicate NBs (magenta) and progeny (cyan), enlarged in the top right insets; a dashed white line indicates the boundary to the optic lobe. **C’**) A typical individual dividing NB from a confocal time-lapse image sequence of the brain lobe. The NB is outlined (dashed white line) and indicated with a magenta arrowhead, the progeny (GMC) is indicated by a cyan arrowhead. **C”**) Plot of NB division rate for cultured L3 brains shows that division rate of NB does not significantly decrease over at least 22 h under imaging conditions (ns, one-way ANOVA), calculated from measured cell cycle lengths. **D’**) Typical GMC division in an intact larval brain. The first row of panels shows production of a GMC (cyan arrowhead) by the dividing NB (magenta arrowhead, dashed white outline). Second row of panels, GMCs are displaced over the next 6 to 8 h by subsequent NB divisions, the path of displacement is indicated by the dashed yellow arrow. The last two panels (10 to 18 min) show the division of a GMC (green arrowhead, progeny yellow arrowheads). **D”**) Plot showing the rate of GMC division in the *ex vivo* brain does not change with time in culture (ns, one-way ANOVA), calculated from the number of GMC division events in 4 hours. Error bars on plots are standard deviation. Scale bars **B** 50 µm; **C, D** 10 µm. *See also Figure S1*.

To assess whether our culturing and imaging protocol supports normal development, we used a number of criteria. We found that by all the criteria we measured, brain development is normal in our *ex vivo* conditions. First, the cultured *ex vivo* brains do not show signs of damage during preparation, which can be easily identified as holes or lesions in the tissue that expand with time in culture. Second, our cultured larval brains consistently increase in size as they progress through development (Figure S1). Third, using our approach, we recorded average division rates of 0.66 divisions/hour (∼90 min per cycle, Figure 1C) for the Type 1 NB of the central brain (Figure S1 A’), at the wandering third instar larval stage (wL3), as previously published (Homem *et al*., 2013; Bowman *et al*., 2008; Movies S1 and S2). We note here that experiments were performed at 21°C, which differs from some developmental studies performed at 25°C. Type I NBs were identified by location according to Homem & Knoblich (2012). Fourth, we rarely observed excessive lengthening or arrest of the cell cycle in NBs over a 22 h imaging period, which is approximately the length of the wL3 stage (Figure 1C). With longer duration culture and imaging, up to 48 h, we observe an increase in cell cycle length, which might be expected for wL3 brains transitioning to the pupal state (Homem *et al*. 2014). Finally, we observed normal and sustained rates of GMC division throughout the imaging period that correspond to the previously described literature in fixed brain preparations (Bowman *et al*. 2010; Figure 1D; Movie S3). We conclude that our *ex vivo* culture and imaging methods accurately represent development of the *Drosophila* brain and support high time and spatial resolution imaging for quantitation of cell numbers and division rates.

### CytoCensus enables easy automated quantification of cell types in time-lapse movies of developing intact larval brains with modest training

Progress in elucidating the molecular mechanisms of regulated cell proliferation during larval brain development has largely depended on the characterisation and quantification of mutant phenotypes by painstaking manual image analysis (for example, Neumüller *et al*., 2011). However, the sheer volume of image data produced by whole brain imaging experiments means that manual assessment is impractical. Therefore, we attempted to use freely available image analysis tools in an effort to automate the identification of cell types. We found that none of the available off-the-shelf image analysis programs perform adequately on our complex 3D datasets, in terms of ease of use, speed or accuracy (Table 1A). Neuroblast nuclei are large and diffuse, which means that conventional spot detectors (e.g. TrackMate) struggle to identify them. Similarly, image segmentation tools (such as RACE, Ilastik and WEKA) struggle to segment NB marked by microtubule labels as they vary significantly in appearance with the cell cycle and cell boundaries may appear incomplete. To overcome these limitations, we developed CytoCensus, an easily deployed, supervised machine learning-based image analysis software (Figures 2; S2). CytoCensus facilitates automated detection of cell types and quantitative analysis of cell number, distribution and proliferation from time-lapse movies of multichannel 3D image stacks even in complex tissues. A full technical description of the algorithm and User Guide is available in the Supplemental Information.

**Table 1.** Cytocensus out-performs other freely available programs for cell class identification. **A)** Performance assessment for a series of freely available tools in identifying NB from a typical 4D live-imaging time-series of the generic cytological markers Jupiter::GFP / Histone::RFP, expressed in larval brains. Comparison to CytoCensus is made on the same computer, including time taken to provide user annotations for a standard data set (150 or 35 time-points, 30-Z). **B)** Direct comparison of Ilastik vs CytoCensus in automatically identifying cell centres in a crowded 3D data set. To facilitate fair comparison, a “neutral challenge dataset” was used (Main Text). F1 score is intuitively similar to accuracy of detection. Values ± standard deviations are shown, n=25 images. Computer specifications: MacBook Pro11,5; Intel Core i7 2.88GHz; 16GB RAM. For manual annotations, the time taken to annotate the full dataset was estimated from the time to annotate 10 time-points. Values ± standard deviations are shown, n = 3. Fiji, ImageJ V1.51d (Schindelin *et al*. 2012; FIJI, local threshold V1.16.4 (http://imagej.net/Auto_Local_Threshold); FIJI-WEKA, WEKA 3.2.1 (Arganda-Carreras *et al*., 2016); RACE (Stegmaier *et al*. 2016); TrackMate (Tinevez *et al*. 2016); Ilastik (V1.17) (Logan *et al*., 2016; Sommer *et al*., 2011).

|  | Manual | Fiji/auto<br>local<br>threshold | TrackMate<br>Spot<br>Detection | RACE | Ilastik<br>(1.17) | Fiji<br>WEKA | CytoCensus<br>V0.1 |
| --- | --- | --- | --- | --- | --- | --- | --- |
| **Total Parameters**<br>**to select** | - | **1** | **4** | 8 | 67<br>(48) | 25 | 6 |
| **Handles 4-D**<br>**easily** | - | NO | **YES** | **YES** | **YES** | NO | **YES** |
| **Time to Train**<br>**model (min.)** | - | N/A | N/A | N/A | 15 | 18 | **6** |
| **Time to Run**<br>**(min. including**<br>**postprocessing)** | 550<br>(equivalent) | **5** | **1** | 16 | 70 | 105 | 19 |
| **F1-score** | - | FAIL | 0.11<br>$\pm 0.09$ | 0.17<br>$\pm 0.01$ | **0.76**<br>**$\pm 0.01$** | 0.62<br>$\pm 0.07$ | **0.96**<br>**$\pm 0.01$** |

**Table 1. Cytocensus out-performs other freely available programs for cell class identification.**
|  | Ilastik (1.17)<br>(raw) | Ilastik<br>(1.17)<br>(post-processed) | CytoCensus<br>V0.1 |
| --- | --- | --- | --- |
| **CPU time (hours)** | 82 | 83 | **12** |
| **Precision**<br>**(True Positive Rate)** | 0.39±0.19 | 0.86±0.10 | **0.98±0.05** |
| **Recall**<br>**(Positive Predictive Value)** | 0.15±0.10 | 0.90±0.07 | **0.98±0.05** |
| **F1-score (max =1.0)** | 0.21±0.13 | 0.88±0.09 | **0.98±0.05** |

**Figure 2.**
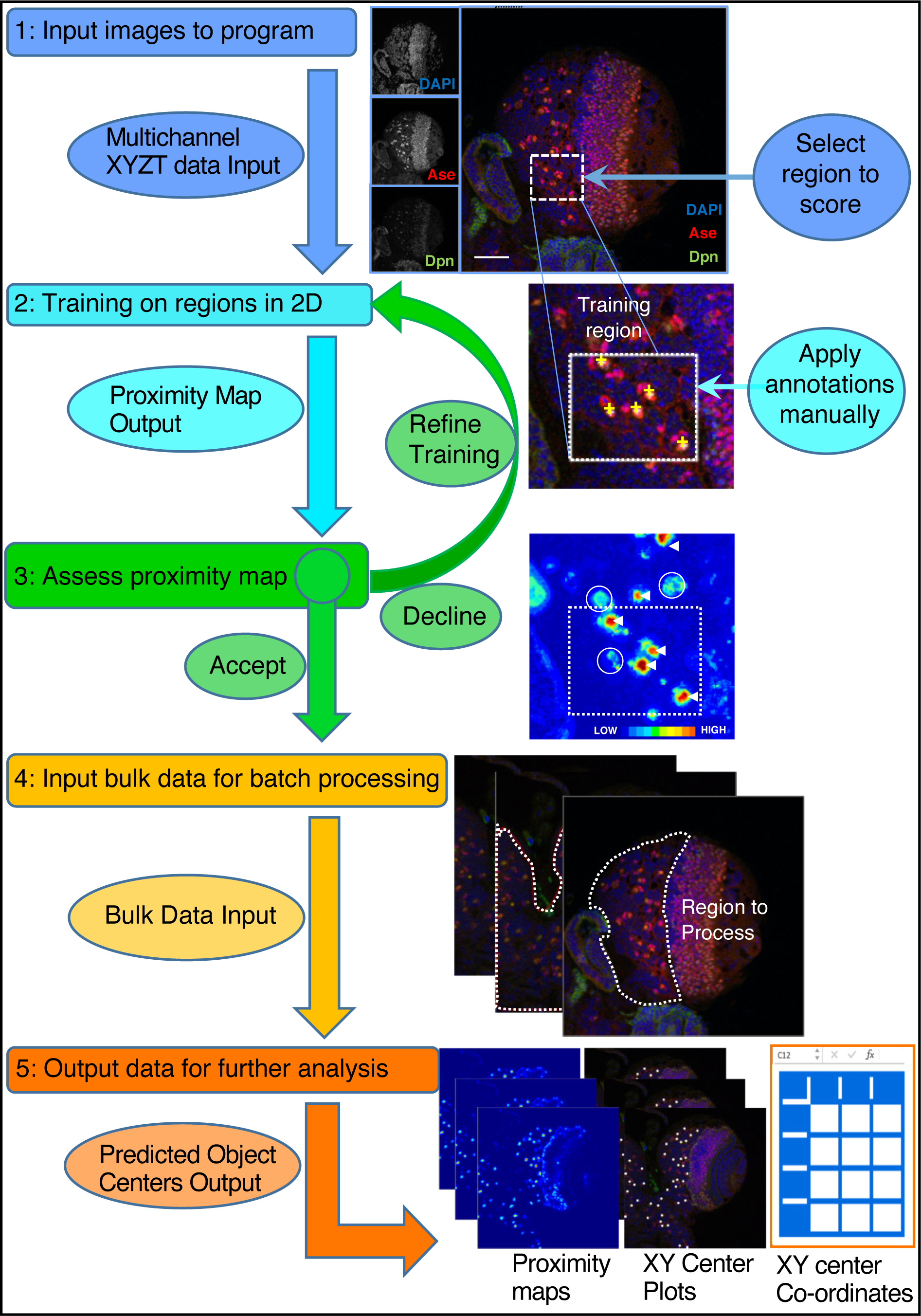
CytoCensus analysis workflow. Refer to the Main Text and Materials & Methods for details. Training is performed by single click annotation (yellow crosses) within a user defined region of interest (ROI, white dashed square) to identify the cell class of interest. The resultant proximity map for cell class identification (∼probability score for object centres) is evaluated manually to assess the success of training (white arrows indicate good detections and circles indicate where more training may be required). A successful identification regime (Model) is saved and may be used to batch process multiple image data sets. Multiple outputs are produced including a list of the co-ordinates of identified cells. Multiple identification regimes can be sequentially applied to identify multiple cell classes from a single data set. *See also Figure S2*.

To optimise its effectiveness, we developed CytoCensus with a minimal requirement for supervision during the training process. We developed an implementation of supervised machine learning (see Supplemental Information), in which the user trains the program in 2D on a limited number of images (Figure 2). In this approach the user simply selects, with a single mouse click, the approximate centres of all examples of a particular cell type within small user-defined regions of interest in the image. This makes CytoCensus is more convenient and faster than other machine learning-based approaches, such as FIJI-WEKA (Arganda-Carreras *et al*., 2016) or Ilastik, (Sommer *et al*., 2011), which require relatively extensive and time consuming annotation of the cells by their boundaries. However, this simple training regime requires assumptions of roundness, which precludes direct analysis of cell shape. We explore the extent of this limitation in subsequent sections.

To further optimise the training, our training workflow outputs a “proximity” map. One may think of the proximity map as a probability of how likely it is that a given pixel is at the center of one of the cells of interest. Using this proximity map the user can assess the accuracy of the prediction and, if necessary, provide additional training (Figure 2). This proximity map and the predicted locations of cell centres across the entire volume and time-series are saved and may be conveniently passed to ImageJ (FIJI), or other programs (Schindelin *et al*. 2012) for further processing (Figures 2; S2; S3). After this initial phase of manual user training, the subsequent processing of new unseen data is automated and highly scalable to large image data sets without any further manual user training. To determine the required training, the impact of training level (number of regions used in the training) was assessed on live imaging data sets (See Supplemental Information). The results show that detection accuracy was optimised even with a modest levels of training (Figure S4).

### Cytocensus is a significant advance in automated cell detection in challenging data sets

We assessed the performance of CytoCensus at cell identification on challenging live imaging data sets that were manually annotated by a user to generate “ground-truth” results. Before comparison between applications, algorithm parameters were optimised for the different approaches to prevent overfitting (see Supplemental Information). In our tests we found that CytoCensus outperformed the machine learning based approaches Fiji-WEKA (p=0.005, t-test, n=3) and Ilastik (p=0.007, t-test, n=3), and other freely available approaches, in the accuracy of NB detection, speed and simplicity of use (Figure 3A; Table 1A). We calculated a metric of performance, intuitively similar to accuracy, which is known as the F1-score, with a maximum value of 1.0 (see Supplemental Information; Table 1A). We found that the best performing approaches on our complex datasets were Ilastik and CytoCensus, which are machine learning based. It is likely that both approaches might be further improved with additional bespoke analysis, specific to each data set, however this would limit their flexibility and ease of use.

**Figure 3.**
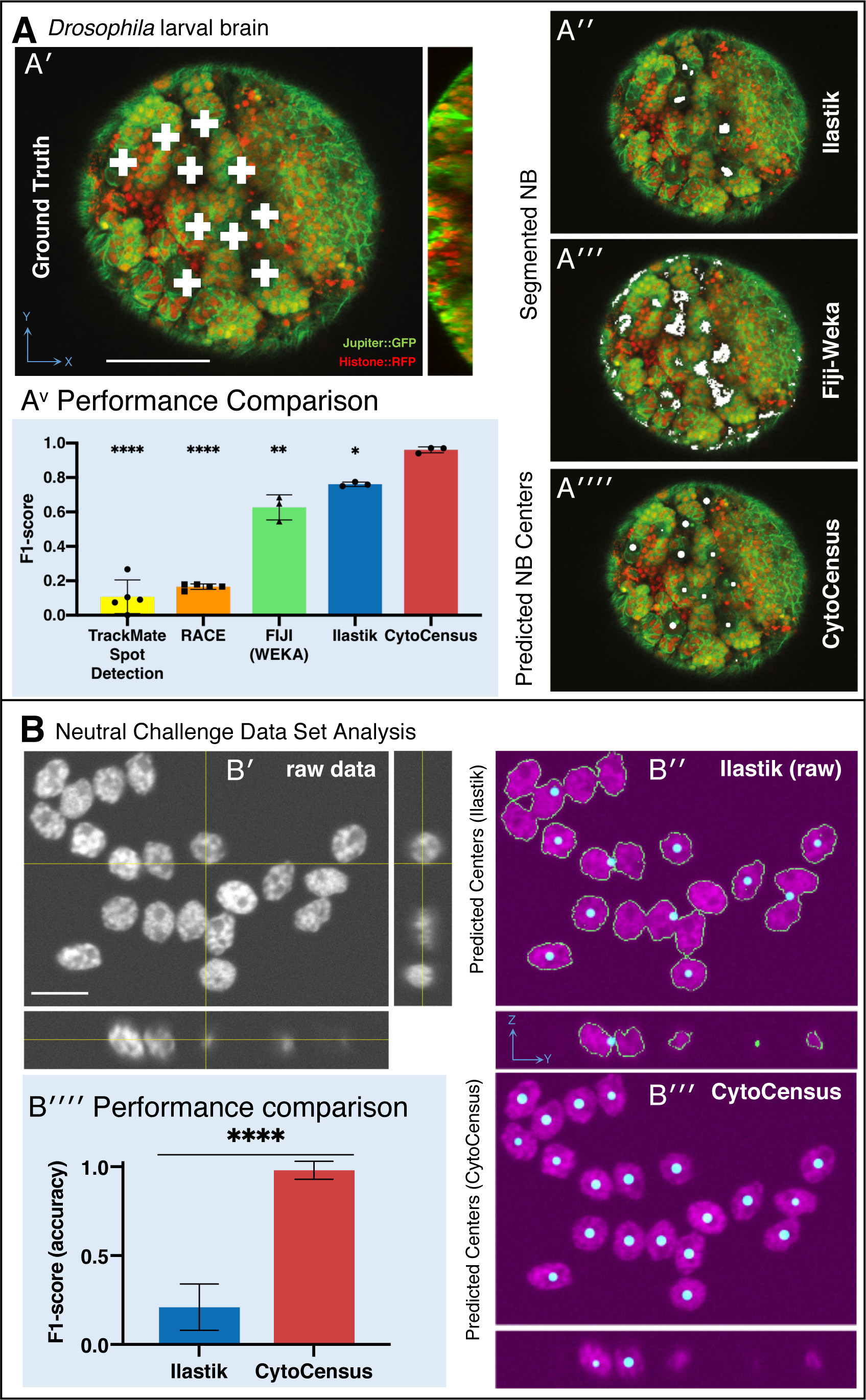
Validation of CytoCensus performance. **A**) Performance in identifying NB from 3D confocal image data of a live brain labelled with Jupiter::GFP, Histone::RFP. **A’**) Ground Truth manual identification of NB centres. **A”** to **””**) Output images comparing NB identification by Ilastik, Fiji-Weka and CytoCensus, white overlay. **A**^**v**^) Plot comparing object centre detection by TrackMate spot detection, RACE, Fiji-Weka, Ilastik and CytoCensus (error bars are standard deviation). CytoCensus achieves a significantly better F1-score than Ilastik (p=0.01, n=3) and FIJI (p=0.005, n=3). (one-way RM-ANOVA with post hoc t-tests) **B**) Comparison of algorithm performance for a 3D neutral challenge data set (**B’**, see Supplemental Information). **B”, B”’**) Output images comparing object centre determination by Ilastik and CytoCensus. Segmentation results are shown as green outlines, object centre determination is show as a cyan point. **B””**) Plot comparing object centre determination accuracy for the 3D neutral challenge dataset (error bars are standard deviation; p<=0.0001, Welch’s t-test, n=25). Scale bars **B** 20 µm; **A’** 50 µm. *See also Figures S3, S4*.

To further critically assess the performance of CytoCensus, we used an artificially generated “neutral challenge” 3D dataset, which facilitates fair comparison (Figure 3B). We used a dataset of 30 images of highly clustered synthetic cells, in 3D, with a low signal to noise ratio (SNR), obtained from the Broad Bioimage Benchmark Collection (see Supplemental Information). We selected this dataset because it has similar characteristics to our live imaging data. Using this dataset we directly compared the abilities of Ilastik (Figure 3B”) and CytoCensus (Figure 3B”’), to identify cell centres in 3D. In both cases we trained on a single image, optimised parameters on 5 images, and evaluated performance on the remaining 25 images. We found that CytoCensus (Table 1B, F1-score: 0.98±0.05) outperforms Ilastik (Table 1B, F1-score: 0.21±0.13) in the accuracy of cell centre detection (Figure 3B) even after the Ilastik results were post-processed to aid separation of touching objects (Table 1B, revised F1-score: 0.88±0.09). We conclude that CytoCensus is significantly more accurate than Ilastik at identifying cells when both are tested out-of-the-box on neutral challenge data (Figure 3B”” p<=0.0001, Welch’s t-test, n=25).

We conclude that CytoCensus represents a significant advance over the other current freely available methods of analysis, both in ease of use and in ability to accurately and automatically analyse cells of interest in the large volumes of data resulting from live imaging of an intact complex tissue such as a brain. This will greatly facilitate the future study of subtle or complex mutant developmental phenotypes.

### Using CytoCensus to analyse the over-growth phenotype of *syncrip* knockdown larval brains

To demonstrate the power and versatility of using CytoCensus in the analysis of a complex brain mutant phenotype, we characterised the brain overgrowth phenotype of *syncrip* (*syp*) knockdown larvae (Figure 4A). SYNCRIP/hnRNPQ, the mammalian homologue of Syp, is a component of RNA granules in the dendrites of mammalian hippocampal neurons (Bannai, H., *et al*., 2004). Syp also determines neuronal fate in the *Drosophila* brain (Ren *et al*., 2017; Liu *et al*., 2015), NB termination in the pupa (Yang *et al*., 2017) and is required for neuromuscular junction development and function (McDermott *et al*., 2014; Halstead *et al*., 2014). *syp* has previously been identified in a screen for genes required for normal brain development (Neumüller *et al*., 2011), although the defect was not characterised in detail.

**Figure 4.**
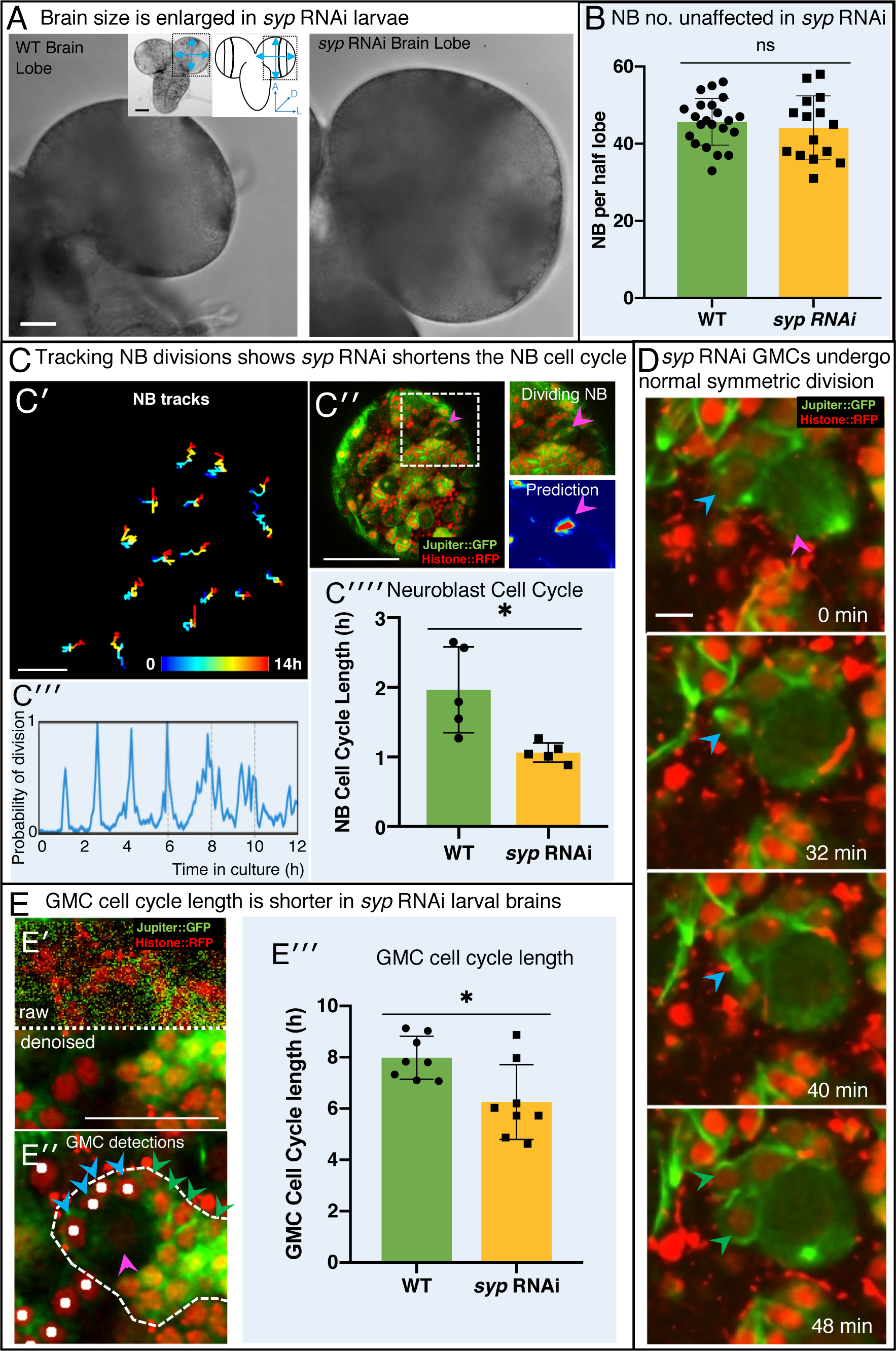
Knockdown of Syncrip protein in NB causes larval brain enlargement. **A)** Brightfield images of freshly isolated brains from third instar WT (OregonR) and *syp* RNAi larvae, respectively. Inserts in (**A**) show the region of the brain imaged and the measurements taken to compare brain size. **B**) Chart comparing NB numbers showing that *syp* RNAi knockdown does not have a significant effect on NB number/brain (ns, t-test, WT n = 22; RNAi n = 15). NB were identified by Dpn labelling and the average count for a comparable volume of a single optic lobe CB region is shown. **C)** Automated identification of NB division using CytoCensus: **C’)** Tracking of NB centres, based on CytoCensus detections, over 14 hours; **C”**) raw image showing single timepoint from live, 3D time-lapse, confocal imaging (insert = single dividing NB, showing CytoCensus prediction of a dividing NB); **C”’)** graph of division of a single tracked NB over 14h; **C””**) average NB (6-9 NB/brain) cell cycle length is reduced in *syp* RNAi knockdown brains (p=0.02, Welch’s t-test, n=5 brains). **D**) Sequence of confocal images from a typical 3D time-lapse movie showing that in *syp* RNAi brains, GMCs divide normally to produce two equal sized progeny that do not divide further. **E**) Semi-automated analysis of GMC division by CytoCensus shows that GMC cell cycle length is reduced in *syp* RNAi brains. **E’)** Single image plane taken from a 3D time-lapse, confocal image data set (imaged at one Z-stack / 2 min). showing raw image data (top) and denoised (bottom). **E”**) CytoCensus GMC detections (cyan) with a single NB (magenta), and NB niche (dotted white line), shows GMCs are detected but neurons (green) are not. **E”’**) Plot of GMC cell cycle length, which is decreased in *syp* RNAi brains compared to WT (p=0.01, Welch’s t-test, n=8 GMCs from 3 brains). Scale bars in **A** 50 µm; **C’** 20 µm; **C”** 50 µm; **D** 5 µm; **E** 25 µm. *See also Figure S5, S6*.

In light of these studies, we wanted to understand the defect caused by Syp on brain development in more detail. We therefore examined *syp* -/- brains (eliminating Syp expression in the NB lineages) and found that in early wL3, brains were significantly enlarged compared to WT larvae at the same stage of development (p<0.0001, t-test, Figures 4A, S5A). *syp* brain lobes exhibit a 23% increase in diameter (WT 206.5 µm ± 5.0, n = 10, *syp* 253.7 µm ± 11.0, n = 5), and a 35% increase in central brain (CB) volume. Significantly, a more specific RNAi knockdown of *syp* driven under the *inscuteable* promoter, which is expressed primarily in NB, and GMCs, demonstrates a similar increase in CB diameter (p=0.002, 13% larger than WT; 234 µm ± 17.0, n=12; Figure S5A). Our data begs the question as to how the removal of *syp* from the neural lineages causes such a significant increase in central brain size.

We tested whether this brain overgrowth is caused by additional ectopic NBs, as has been previously described for other mutants (Bello *et al*., 2006). We used CytoCensus to accurately determine the total number of NBs in the CB of fixed *syp* knockdown verses WT wL3 brains. Our results show that wL3 brains with *syp* RNAi knockdown have no significant difference in ventral NB number compared to WT (Figure 4B; WT 45.6 ± 1.3, n = 22, *syp* RNAi 44.1 ± 2.1, n = 15). We conclude that a change in NB number is not the underlying cause of brain enlargement observed in *syp* RNAi and hypothesise that a change in NB division rate or that of their progeny might be responsible.

### *syp* RNAi knockdown brains exhibit an increased NB division rate

To investigate whether an increase in NB division rate contributes to the brain overgrowth observed in *syp* knockdown larvae, we examined the rate of NB division in living brains using our optimised culturing and imaging methods, followed by CytoCensus detection and tracking.

First we perform 3D NB detections using CytoCensus (as shown previously in Figure 3A), and we fed this input into TrackMate, a simple tracking algorithm. Without the CytoCensus detections, TrackMate spot detection performs poorly on the raw data (F1 score 0.11+0.09), and tracking is all but impossible. Applying TrackMate to the proximity maps generated by CytoCensus dramatically improves TrackMate detections (F1 score 0.92+0.02, Figure S6A). As a result, 16 out of 17 NB were successfully and accurately tracked for over 20 h in our tests (Figure 4C’,Figure S6AV).

In order to follow the NB cell cycle, we next showed CytoCensus can accurately identify individual dividing NBs in live image series, both in WT (Figure 4C”) and in RNAi brains (Figure S6B). We detected dividing NB by training on NB with visible spindles using CytoCensus, and used this output to create plots of division for each NB (Figures 4C”’, S6C-D). Using these plots, we measured the cell cycle length of NBs in wild type and *syp* RNAi brains and found that, on average, *syp* RNAi NBs have a 1.78-fold shorter cell cycle compared to WT (p=0.02, Welch’s t-test, N=5 brains; Fig 4C””). We propose that this shorter cell cycle length (i.e. an increased division rate) in the *syp* knockdown is the primary cause of its increased brain size. These results illustrate the potential of CytoCensus to analyse the patterns of cell division in a complex, dense tissue, live, in much more detail than conventional methods in fixed material.

### GMC cell cycle length is slightly decreased in *syp* RNAi brains

We also investigated GMC behaviour in the CB region of *syp* RNAi and WT larval brains, to test whether an aberrant behaviour of mutant GMCs could also contribute to a brain enlargement phenotype. Given that GMCs are morphologically indistinguishable from their immature neuronal progeny (which makes them particularly difficult to assess) we had to identify GMCs by tracking them from their birth in a NB division to their own division into two neurons. To achieve this goal required us to use high temporal resolution imaging and patch based denoising (Materials & Methods) which allowed us to confirm that normal, symmetric GMC divisions occurred with the correct timing and resulted in two daughter cells (which did not regrow or divide further), both in WT and *syp* RNAi (Figure 4D).

Using our refined culture and imaging conditions, we trained CytoCensus to successfully detect GMCs in denoised images (Figure 4E’-E”) and, similarly to NB, track them with a trackpy based script (See Materials & Methods and Supplemental Information). Unlike in the case of NB tracking, GMCs do not go through repeated cycles of division, so following automated detection, for each GMC, we manually identified the birth and final division and additionally corrected any tracking errors. This semi-automated tracking allowed us to compare the cell cycle length of GMCs in multiple brains over 12h time-lapse movies for the first time (Figure 4E”’). In *syp* RNAi, we find a small but significant shortening (p=0.01, Welch’s t-test) of the cell cycle compared to WT (8.00h +/− 0.89, n=8 WT; 6.25h +/− 1.45, n=8 *syp* RNAi). However, while we conclude that GMC cell cycle length is decreased by 20%, GMCs terminally divide normally (representative example, Figure 4D), and we see no evidence of further divisions in the neurons. We take this to mean that no additional cells are produced by GMC or neuron division and therefore brain size is not significantly affected. We conclude that the cause of the enlarged brain size in *syp* RNAi brains is an increase in NB division rate resulting in more GMCs and their progeny than in WT.

### NB division rate is consistently heterogenous in *Drosophila* brains

Most current methods for measuring NB division rates produce an average rate for whole brains rather than providing division rates for individual NBs. It has previously been shown that NB lineages give rise to highly variable clone size (30-150 neurons for Type I neuroblasts). The origin of this diversity has primarily been attributed to patterned cell death (Yu *et al*., 2014), but the importance of NB division rate in determining clone size is less well understood. Using live imaging and CytoCensus, however, we were able to quantitate the behaviour of multiple individual NBs over time within the same brain to investigate whether cell division rates are constant or variable across the population. Interestingly, we found that each NB has a constant cell cycle period (Figure 5A), matching observations *in vitro* (Homem *et. al*., 2013). However, there is considerable variation in cell cycle length between NBs within the same brain lobe, (Figure 5A). Given the scale of this variation, which covers more than 2-fold difference in rate, we expect that the regulation of NB division rate is a key factor that contributes to the observed variation in NB lineage size. By comparing the distribution of division rates in individual WT and *syp* RNAi brains, we found that *syp* knockdown NBs have a more consistent division rate in individual NBs (Figure 5B) and between brains (Figure 4C”’), which suggests a role for *syp* in the regulation of NB division rate. Future work using CytoCensus and live imaging would allow one to explicitly link individual NB division rates to atlases of neural lineages and explain the contribution of division rate to each neural lineage.

**Figure 5.**
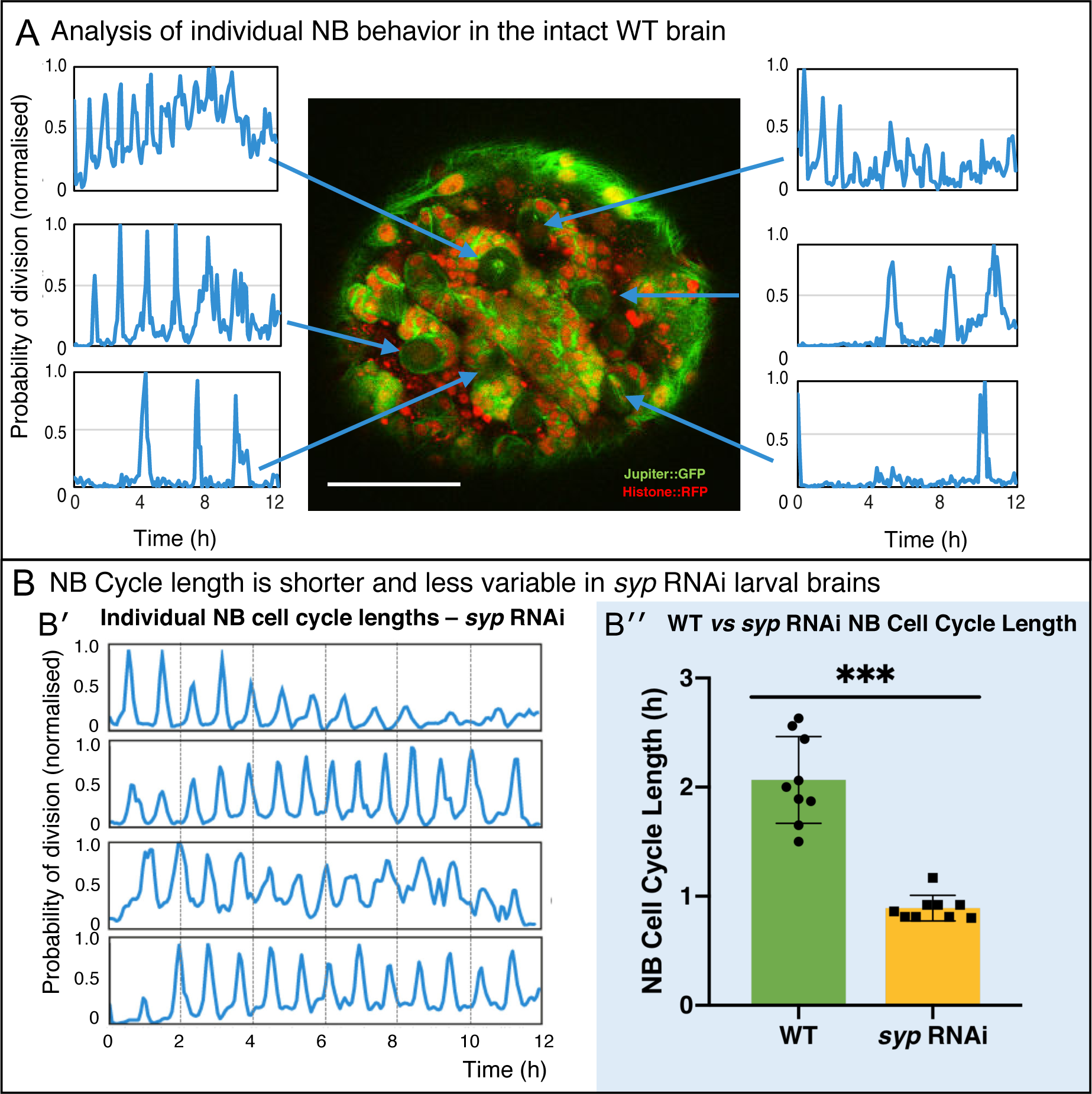
Direct analysis of NB division from time-lapse imaging of live explanted larval brains. **A**) Using the proximity map output of CytoCensus, individual NBs can be followed through their cell cycle. Arrows: Individual NB locations, and the corresponding proximity map output plotted over time for that NB. **B)** Comparison of WT and *syp* RNAi NB: **B’**) analysis of cell cycle over time for individual NBs from a *syp* RNAi brain; **B”**) comparison of cell cycle lengths for individual NB in a single WT vs *syp* RNAi brain (p<0.001, Welch’s t-test, n=9). Scale bar 40 µm. *See also Figure S6*.

We conclude that analysing live imaging data with CytoCensus can provide biological insights into developmental processes that would be difficult to obtain by other means. However it was important to establish the use of CytoCensus in other situations outside *Drosophila* tissues, especially in vertebrate models of development.

### Directly quantifying cell numbers enhances the analysis of zebrafish retinal organoid assembly

To test the utility of CytoCensus for the analysis of complex vertebrate tissue, we first analysed Zebrafish tissue, an outstanding model for studying development with many powerful tools, such as the Spectrum of Fates (SoFa) approach (Almeida *et al*., 2014), which marks cells from different layers of the Zebrafish retina by expression of distinct fluorescent protein labels. Previously published work by Eldred *et al*. (2017) studying eye development in artificial Zebrafish organoids, provided an excellent example of material that was previously analysed using bespoke MATLAB image analysis software that measured only the cumulative fluorescence at different radii from the organoid centre. While this was sufficient for a summary of organoid organisation, future research will require the ability to examine organoids at the single cell level, particularly in cases where layers are formed from a mixture of cell types or cell types are defined by combinations of markers. We deployed CytoCensus to this end, without the need for bespoke image analysis, in directly locating and counting cells (Figure 6A).

**Figure 6.**
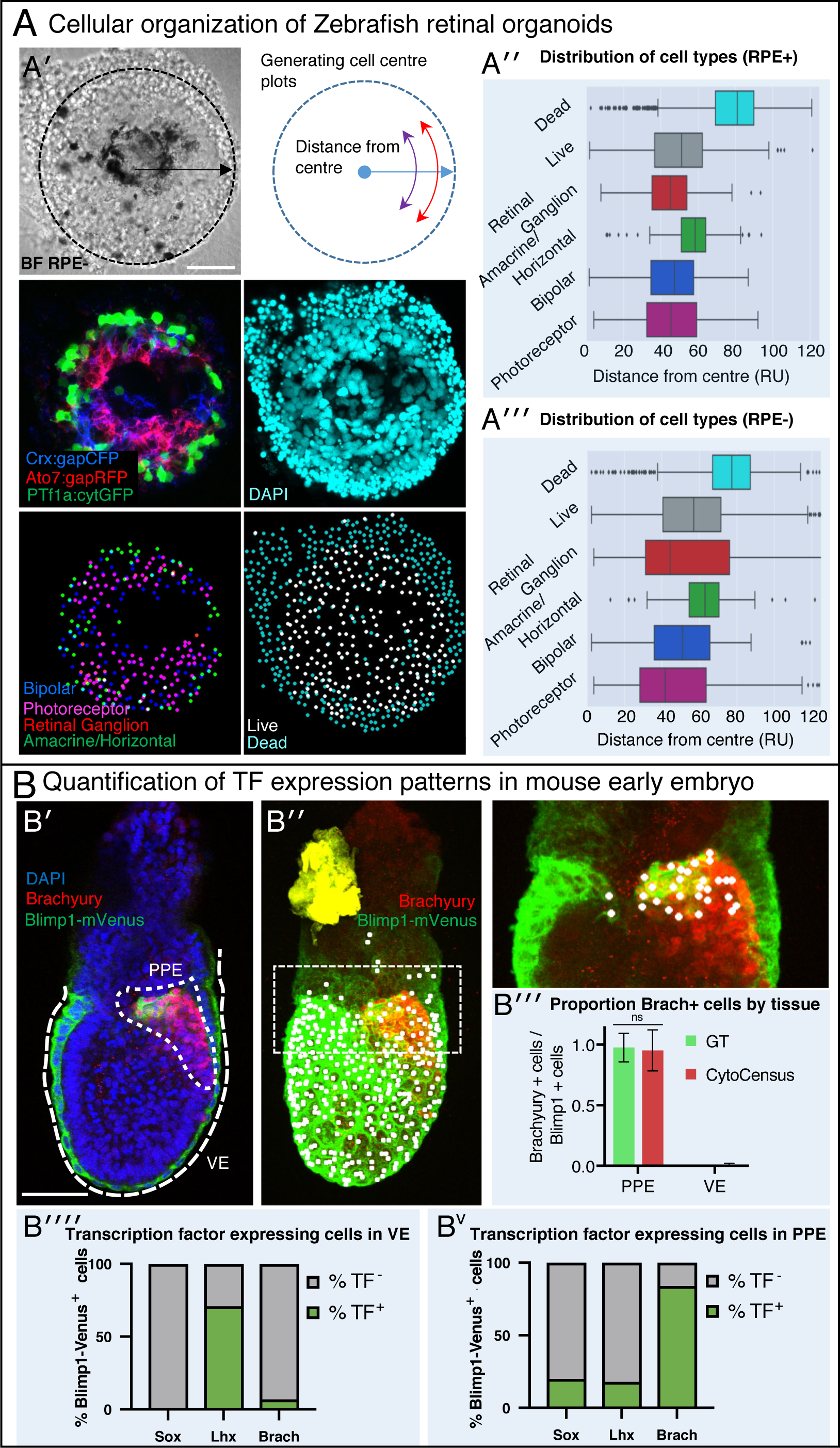
A generally applicable automated analysis tool to assess tissue development. **A**) Automated analysis of Zebrafish retinal organoids at the single cell level (Raw data from Eldred *et al*., 2017). **A’** Top: brightfield image and diagram indicating the location of cells was defined as displacement from the organoid center. Middle: Cell fate marker expression (Crx:gapCFP; Ato7:gapRFP; PTf1a:cytGFP) and DAPI. Bottom: Cell centre identification by CytoCensus for the different cell types as defined by the labelling profiles (Bipolar, Photoreceptor, Retinal Ganglion, Amacrine/Horizontal, Live/Dead). (**A”-A”’**) Radial distribution of the different cell types determined from cell centre identifications by CytoCensus; the effects on organoid organisation of the presence (**A”**) or absence (**A”’**) of retinal pigment epithelium (RPE) cells is examined (ns, one-way ANOVA). RU = Radial Units, normalised to a radius of 100 (see Materials & Methods) **B**) Automated quantification of TF expressing cells in a fixed early streak stage mouse embryo (e6.5) labelled for transcription factors, Blimp1-mVenus and DAPI. **B’**) A medial confocal section showing Brachyury in the primitive streak in the proximal posterior epiblast (PPE) and visceral endoderm (VE, highlighted cortical tracing). **B”**) Cortical image of the same mouse embryo overlaid with total cell centre predictions by CytoCensus of Brachyury positive cells; insert to the right is a zoomed in image of the highlighted rectangle showing only cell centre predictions in a single medial plane. **B”’**) Comparison of CytoCensus and manual Ground Truth (GT) measurements of the proportion of Brachyury positive cells from 2D planes in the VE and PPE (ns, t-test, n=3). **B””-Bv**) Proportion of transcription factor positive cells (TF) in, using CytoCensus measurements in 3D according to tissue regions (PPE and VE) defined in (**B’**). Scale bars 25 µm in **A**; 100 µm in **B’**. *See also Figure S7*.

Using CytoCensus, we trained multiple models on subsets of the raw data (Figure 6A’, gift from the William Harris lab), corresponding to each of the different cell types. Applying our models to the remainder of the dataset, CytoCensus was able to identify individual cells (Figure 6A, bottom panels), allowing an analysis of cellular distribution that would not be possible from cumulative fluorescence measurements. We then calculated the number of cells found at different distances from the center of the organoid (Materials & Methods, Figure 6A”-A”’). Using this approach, we reproduced the previously published analysis (Eldred *et al*., 2017), mapping the different cell distributions in the presence and absence of retinal pigment epithelium cells. We show that CytoCensus produces similar results to Figure 2 of Eldred *et al*. (2017), but with identification of individual cells and without the need for a dedicated image analysis pipeline (Figure 6A”-A”’). In particular, we are able to produce an estimate of the distribution of the photoreceptor (PR) cell class, which is defined by a combination of markers (Crx::gapCFP, Ato7::gapRFP) that could not be separated from other cell types in the original analysis.

Given that the SoFa markers support the study of live organoid development, and CytoCensus can be used to identify cells based on the SoFa markers, we expect CytoCensus could easily be used to analyse live organoid development along similar lines to our *Drosophila* analysis. We conclude that CytoCensus is an effective tool to investigate the distribution of cell types in the assembling retinal organoid, with the potential to analyse other complex Zebrafish tissues.

### CytoCensus facilitates rigorous quantification of TF expression patterns in mouse embryos

Mouse models are widely used to understand developmental processes in the early embryo. In such work, genetic studies have been fundamental in understanding the molecular mechanisms underlying important lineage decisions (Piliszek *et al*., 2016; Arnold & Robertson, 2009). However, assessment of changes in cell numbers and distribution frequently relies on manual counting and qualitative estimation of phenotypes. We tested the ability of CytoCensus to provide quantitative data on the number of transcription factor positive cells in the early post-implantation embryo for each of the transcription factors Brachyury, Lhx1 and Sox2. Using CytoCensus we quantitated the number of cells that express each of these transcription factors in two regions of interest: the visceral endoderm (VE) and the proximal posterior epiblast (PPE), where primordial germ cells (PGCs) are specified. We also analysed the distribution of Blimp1-mVenus in membranes in both the VE and PGCs (Ohinata *et al*., 2008; Figure 6B’, B”).

Using CytoCensus we identified all Blimp1 expressing cells and mapped them to structures of interest using a 3D ROI (Figure 6B’ marked regions). We then used CytoCensus to identify cells expressing both Blimp1 and Brachyury in the proximal posterior epiblast (PPE) (Figure 6B” and insert). We note that CytoCensus could be used to successfully detect cells of the VE and PGCs, despite the fact that they are frequently far from round. CytoCensus is able to detect these cells, almost as well as truly round cells, by integrating information from the nuclear and membrane markers to produce robust cell centre detections. Our analysis highlights the enrichment of Brachyury in the developing PGCs and their almost complete absence from the VE, which matches well with manual 2D quantification (Figure 6B”’). Repeating this analysis for the transcription factors Sox2 and Lhx1 highlights a differential expression of the transcription factors (Figure 6B””-V). These proportions match well with qualitatively reported expression patterns in the field (Piliszek *et al*., 2016). Our results demonstrate how CytoCensus can be used to produce a robust and detailed quantitation of cell type and TF expression in specific complex mouse tissues using standard markers, improving on the standard qualitative analysis.

Taking our results in their entirety, in *Drosophila*, Zebrafish and mouse, we illustrate the wide applicability of CytoCensus to transform the quantitative analysis of any complex tissue. CytoCensus makes it possible without bespoke programming to quantitate cell numbers and their divisions in complex living or fixed tissues in 3D.

## Discussion and Limitations

Progress in understanding the development and function of complex tissues and organs has been limited by the lack of effective ways to image cells in their native context over extended developmentally relevant timescales. Furthermore, a major hurdle has been the difficulty of automatically analysing the resulting large 4D image series. Here, we describe our development of culturing and imaging methods that support long term high resolution imaging of all the cells in intact living explanted *Drosophila* larval brains. This progress relies on optimised dissection and mounting protocols, a simplified culture medium for extending brain viability and the use of patch-based denoising algorithms to allow high resolution imaging at a tenth of the normal illumination intensity. We next describe our development of CytoCensus: a convenient and rapid image analysis software employing a supervised machine learning algorithm. CytoCensus was developed to identify neural stem cells and other cell types, both in order to quantitate their numbers and distribution and to enable analysis of the rate of division on an individual cell level, from complex 3D and 4D images of cellular landscapes. We demonstrate the general utility of CytoCensus in a variety of different tissues and organs.

To image all the cells in an explanted brain, we used very bright generic markers of cellular morphology, which offer major advantages over specific markers of cell identity, as they are more abundant and brighter, allowing the use of low laser power to maximise viability. Markers of cell morphology can also be used in almost all mutant backgrounds in model organisms, unlike specific markers of cell identity, whose expression is often critically altered in mutant backgrounds. However, imaging all the cells in a tissue or organ with generic markers leads to complex images, in which it is very challenging to segment individual cells using manual or available image analysis tools. In contrast to other approaches, we demonstrate that CytoCensus allows the user to teach the program, using only a few examples, by simply clicking on the cell centres. CytoCensus outperforms, by a significant margin, the other freely available approaches that we tested, so represents a step change in the type and scale of datasets that can be effectively analysed by non-image analysis experts. Crucially, CytoCensus analysis combined with cell tracking in extensive live imaging data allows parameters such as cell cycle length to be determined for individual cells in a complex tissue, rather than conventional methods that provide snapshots or an ensemble view of average cell behaviour.

The image analysis approach we have developed depends critically on the use of “supervision” or training regimes which are, by definition, subjective and user dependent. Supervised machine learning methods (Luengo *et al*., 2017; Arganda-Carreras *et al*., 2017; Logan *et al*., 2016; Chittajallu *et al*., 2015; Sommer *et al*., 2011) require the user to provide training examples by manually identifying (annotating) a variety of cells or objects of interest, often requiring laborious “outlining” of features to achieve optimal results. However, our use of a “point and click” interface (Figure S2), to simplify manual annotation, and proximity map output, makes it quick and easy for a user to train and retrain the programme. Using our approach, a user can rapidly move from initial observations to statistically significant results based upon bulk analysis of data.

We show the value of CytoCensus in three key exemplars. In *Drosophila*, we measure cell cycle lengths *ex vivo* in two key neural cell types, revealing the significant contribution of neuroblast division rate to the *syp* RNAi overgrowth phenotype. In Zebrafish organoids, we illustrate that CytoCensus is generally applicable and compatible with other cell types and live imaging markers. We show it is possible to easily characterise organoid organisation at the cellular level, including analysis of cell type which was not previously quantified (Eldred *et al*. 2017). Finally, we quantify TF expression in images of mouse embryos, illustrating how qualitative phenotypes can be straightforwardly converted into quantitative characterisations, even in epithelial tissue which differs from the typical assumptions of round cells.

A technical limitation of our “point and click” strategy is that the program “assumes” a roughly spherical cell shape. This means that cellular projections, for instance axons and dendrites of neurons, would not be identified, and other programs (e.g. Ilastik, etc.) may be more appropriate to answer specific questions that require knowledge of cell shape or extensions. However, we find that the robustness of the CytoCensus cell centres, even with irregular or extended cells can be a useful starting point for further analysis. To this end we configured the output data from CytoCensus to be compatible with other programs, such as FIJI (ImageJ), allowing a user to benefit from the many powerful plug in extensions available to facilitate further extraction of information for defined cell populations from bulk datasets.

With the increased availability of high throughput imaging, there is a greater unmet need for automated analysis methods. Ideally, unsupervised methods will remove the need for manual annotation of datasets, but at present, the tools required are in their infancy. In this context, methods that require minimal supervision, such as CytoCensus are desirable. Machine learning approaches, such as CytoCensus, offer the potential to analyse larger datasets, with statistically significant numbers of replicates, and in more complex situations, without the need for time-consuming comprehensive manual analysis. Easing this rate limiting step will empower researchers to make better use of their data and come to more reliable conclusions. We have demonstrated that analysis of such large live imaging datasets with CytoCensus can provide biological insights into developmental processes in *Drosophila* that would be difficult to obtain by other means, and that CytoCensus has a great potential for the characterisation of complex 4D image data from other tissues and organisms.

## Author Contributions

MH, RMP, LY, TJS, IC, TD and YA designed and performed experiments and MH, RMP, LY, IC, TJS and ID analysed and interpreted data. DW originated the computational approaches used and developed in collaboration with MH the software for 3D analysis in CytoCensus. MH extensively validated and tuned the software to the application of interest and performed analysis of imaging data. LY initiated the project intellectually and the application to Syncrip. DW initiated the computational approaches used. MH, RMP, LY, DW, TJS discussed the results and conclusions, and all authors commented on and contributed to the revision of the manuscript. RMP and ID supervised the project, the biological applications and user interface. RMP, MH and ID wrote the manuscript and all authors contributed to revisions.

## Acknowledgements

We are grateful to: Ivo A. Telley (Instituto Gulbenkian de Ciência) for fly stocks; the Harris and Robertson labs for sharing their imaging data; Jordan Raff and Russel Hamilton for their insightful comments on the results; David Ish-Horowicz and Alfredo Castello for discussions and critical reading of the manuscript. We would also like to thank Tomek Dobrzycki for his contribution to the initial characterisation of the Syp mutant phenotype. Thanks to Andrew Jefferson and MICRON (http://micronoxford.com, supported by a Wellcome Strategic Awards 091911/B/10/Z and 107457/Z/15/Z) for access to equipment and assistance with imaging techniques. This work was supported by: a Clarendon Fellowship (Oxford University Press) to L.Y.; MRC/BBSRC/EPSRC (grant number MR/K01577X/1) and the Wolfson Foundation, Medical Research Council (MRC) Grants MC_UU_12010/Unit Programs G0902418 & MC_UU_12025 supporting D.W.; Wellcome Trust Senior Research Fellowship (081858) to I.D. and supporting R.M.P; Wellcome Trust Four-Year PhD Studentship (105363/Z/14/Z) to T.J.S.; a UKRI MRC grant (MR/S005382/1a, MC_UU_12009) to DW and by funding from the Engineering and Physical Sciences Research Council (EPSRC) and Medical Research Council (MRC) [grant number EP/L016052/1] supporting M.H.

## Materials and Methods

### Fly strains

Stocks were raised on standard cornmeal-agar medium at either 21 °C or 25 °C. To assist in determining larval age, Bromophenol Blue was added at 0.05% final concentration in cornmeal-agar medium. The following *Drosophila* fly strains were used: [Wild-Type Oregon-R]; [Jupiter::GFP;Histone::RFP (recombination on the third)]; [AseGal4>UAS-MCD8-GFP]; [w11180;PBac(PB)sype00286/TM6B]; [Bloomington 9289, w11180 (homozygote syp Null)]; [Df(3R)BSC124/TM6B (crossed to BL 9289 for syp Null)]; [syp RNAi lines - w11180; P{GD9477} v33011, v33012].

### Mouse Embryos

Refer to Simon *et al*., (2017) for details on mouse embryo preparation.

### Fixed Tissue Preparation and labelling

Flies of both genders were raised as described above and larvae from second instar to pre-pupal stages collected and dissected directly into fresh 4% EM grade paraformaldehyde solution (from a 16% stock. Polysciences) in PBS with 0.3% TritonX-100 then incubated for 25 min at room temperature (RT). Following fixation, samples were washed 3 times for 15 min each in 0.3% PBST (1x PBS containing 0.3% Tween) and blocked for 1 h at RT in Immunofluorescence blocking buffer (1% FBS prepared in 0.3% PBST). Samples were incubated with primary antibody prepared in blocking buffer for either 3 h at RT or overnight at 4 °C. Subsequently, samples were washed 3 times for 20 min each with 0.3% PBST followed by incubation with fluorescent labelled secondary antibodies prepared in blocking buffer for 1 h at RT. For nuclear staining, DAPI was included in the second last wash. Samples were mounted in VECTASHIELD (Vector Laboratories) for examination. For details on the preparation and labelling of mouse embryos, refer to Simon *et al*., (2017).

### Culture of live explanted larval brains on the microscope

Brains were dissected from 3rd instar larvae in Schneider’s medium according to https://www.youtube.com/watch?v=9WlIoxxFuy0 and placed inside the wells of a pre-prepared culturing chamber (Figure 1A). To assemble the culturing chamber, 1% low melting point (LMP) agarose (ThermoFischer) was prepared as 1:1 v/v ratio of 1 × PBS and Schneider’s medium (ThermoFisher 21720024) then pipetted onto a 3 cm Petri dish (MatTek) dish and allowed to solidify. After solidification, circular wells were cut out using a glass capillary ∼ 2 mm diameter. To secure the material in place, a 0.5% LMP solution [1% LMP solution brain diluted 1:1 with culturing medium (BCM)] was pipetted into the wells to form a cap. Finally, the whole chamber was flooded with BCM. BCM was prepared by homogenising ten 3rd instar larvae in 200 µl of Schneider’s medium and briefly centrifuge to separate from the larval carcasses. This lysate was added to 10 ml of 80% Schneider’s medium, 20% Foetal Bovine Serum (GibcoTM ThermoFisher), 10 µl of 10 mg/ml insulin (Sigma). A lid is used to reduce evaporation. For GMC imaging we used a solid-agar cap (1-2% LMP agarose) placed directly on top of the brains, which we found was more consistent at holding brains against the coverslip than our earlier approach. We note that care must be taken not to flatten brains during this process, as it appears to result in a higher rate of stalled NB divisions which are likely artefacts. This approach reduced movement in brains significantly, but did not eradicate it - it seems likely remaining movement is the primarily the result of thermal drift of the microscope focus, and is well corrected using image registration.

### Imaging

Confocal, live imaging of *Drosophila* was performed using an inverted Olympus FV3000 six laser line spectral confocal fitted with high sensitivity gallium arsenide phosphide (GaAsP detectors), x30 SI 1.3 NA lens. The confocal pinhole was set to one airy unit to optimise optical sectioning with emission collection. Images were collected at 512×512 pixels using the resonant scanner (pixel size 0.207 µm) and x2 averaging). The total exposure time per Z stack (60) frames was ∼20s. For live culture and imaging the sample was covered with a lid at 21±1°C. Imaging of the GMC cell cycle required increased temporal and spatial resolution, compared to imaging NB: 2 min. time-lapse with 0.2×0.2×0.5 µm resolution. Initial tests indicated that the resulting increased light dosage reduce the number of GMC divisions over time, which we consider to be a sign of phototoxicity. Therefore, we reduced the laser power by approximately a factor of 10 (to ∼12µW at the objective for 488 nm, and 7µW for 561 nm), and used post acquisition patch-based denoising (NDSAFIR, by Kervrann and Boulanger, 2006, implemented as part of PRIISM, with adapt=0, island=4, zt mode and iterations=3 by Carlton *et al*., 2010) to restore image quality. For details on imaging of mouse embryos (Figure 6) refer to Simon *et al*., (2017). Details of organoid imaging can be found in (Eldred *et al*., 2017). Additional live imaging was carried out on a GE Deltavision Core widefield system with a Lumencor 7-line illumination source, Cascade-II EMCCD camera and x30 SI 1.3 NA lens

For imaging of fixed *Drosophila* material, either an Olympus FV1200 or FV1000 confocal was used with x20 0.75 dry or x60 1.4 NA. lenses. Settings were adjusted according to the labelling and were kept consistent within experiments.

For brightfield imaging (Figure S1, S5) a GE Deltavision Core widefield system, Cascade-II EMCCD camera and x30 SI 1.3 NA lens was used. Measurements of brain diameters were performed by hand in OMERO. Reported measurements are the average of one measurement along the longest axis of a brain lobe (passing through the central brain and optic lobe), and another at right angles to that (typically across the medulla).

### Image Analysis (Summary)

All programs used for image analysis were installed on a MacBook Pro11,5; Intel Core i7 2.88GHz;16GB RAM. Basic image handling and processing was carried out in FIJI (ImageJ V1.51d; http://fiji.sc, Schindelin *et al*. 2012). The CytoCensus software, and additional scripts were written in Python, a detailed technical description is given in the Supplemental Information section.

### Data and Software Availability

The following freely available image analysis tools were used: Fiji, ImageJ V1.51d http://fiji.sc, Schindelin *et al*. 2012); Ilastik (V1.17) (http://ilastik.org; Logan *et al*., 2016; Sommer *et al*., 2011). The CytoCensus software can be installed as a stand-alone program: full install available at www.GitHub.com/hailstonem/CytoCensus. Image data was archived in OMERO V5.3.5 (Allan *et al*., 2012; Linkert *et al*., 2010); image conversions were carried out using the BioFormats plugin in Fiji (Linkert *et al*., 2010; https://imagej.net/Bio-Formats).

### Quantification and Statistical Comparison

Mutant comparisons were performed using an appropriate test in GraphPad Prism (see Figure legends for specific tests), typically a Student’s T test, following Shapiro-Wilk test to test normal distribution of the data. A p-value of <0.05 was considered significant. Numbers of replicates typically refer to the number of independent brains and are detailed in the figure legends and main text. Unless otherwise stated, error bars shown are standard deviation.

### Table of Resources

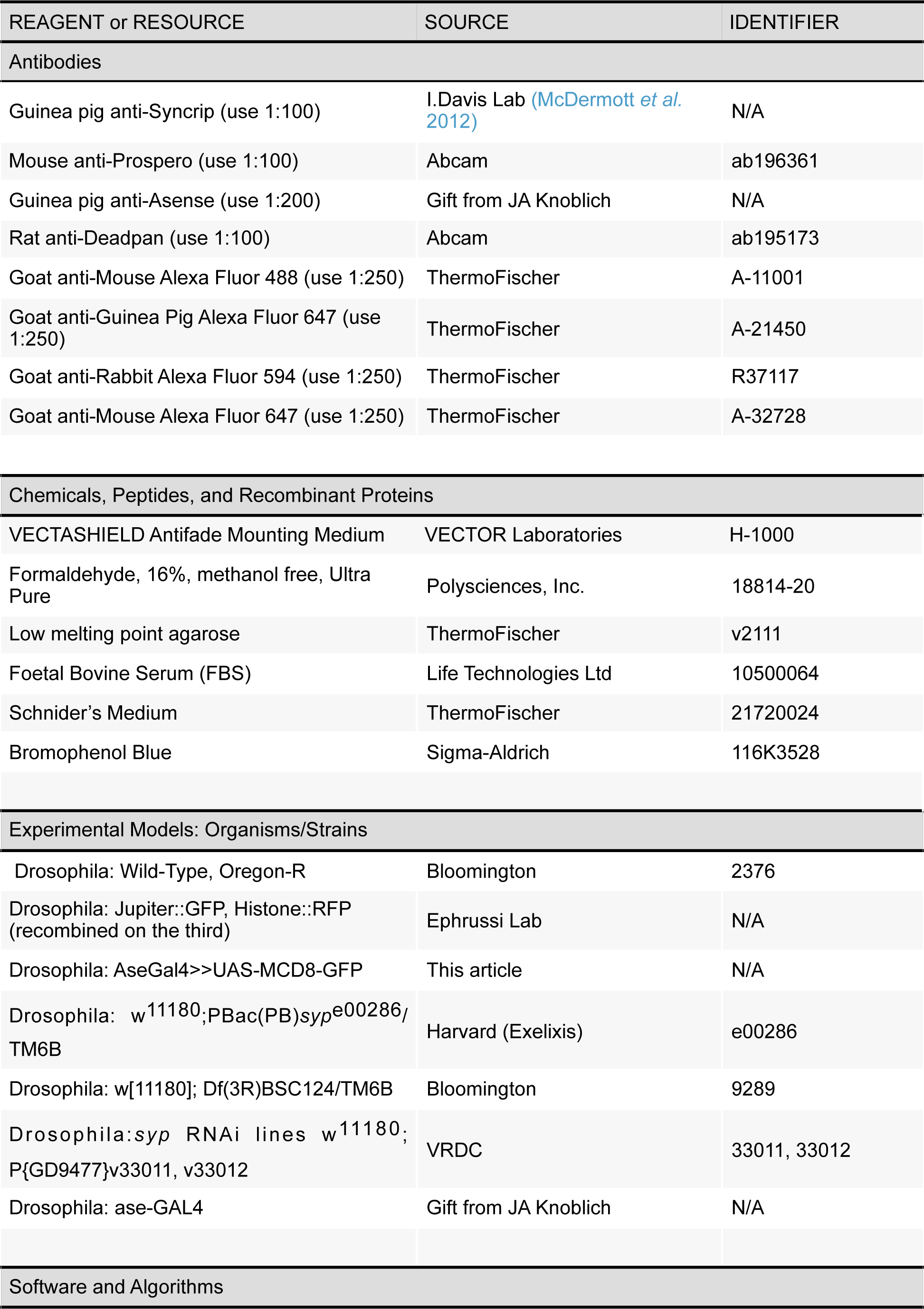
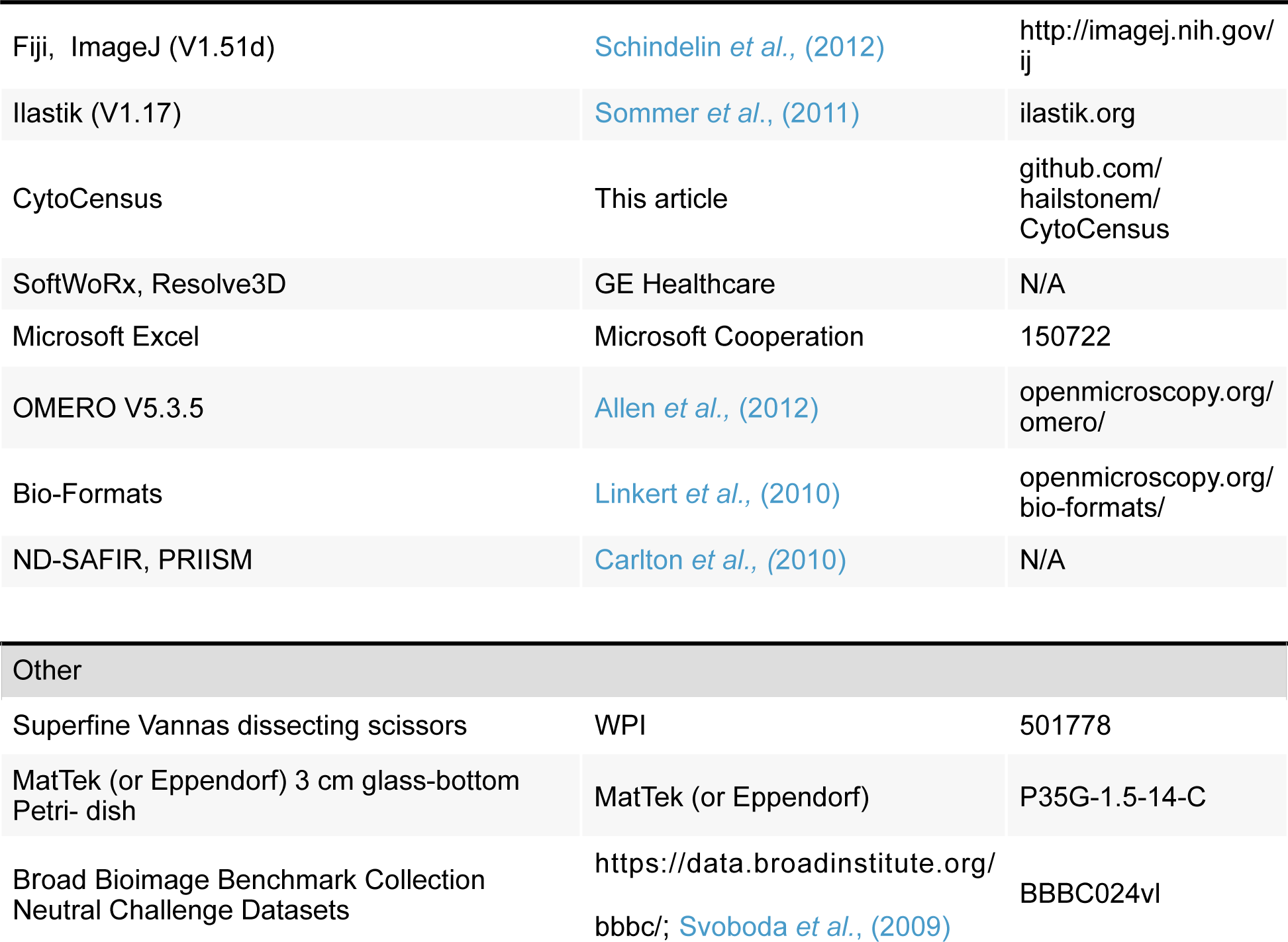

## ADDITIONAL FILES

**Supplemental Movie 1**: (See Figure 1B) Development of a live explanted larval brain under extended time-lapse imaging conditions. Time-series (12h) of one of the brain lobes, collected at 2 minute intervals and displayed at 5 fps. Red: Histone::RFP; Green: Jupiter::GFP. Left: Registered and denoised movie. Right: Raw imaging data. Repeated asymmetric division of a NB regenerates a daughter NB and produces a smaller GMC. Scale bar 10 um.

**Supplemental Movie 2:** (See Figure 1C) Neuroblast division in live explanted larval brains under extended time-lapse imaging conditions. Time-series (13h), collected at 6 minute intervals and displayed at 3 fps. Red: Proximity map for Dividing NB, Blue: Proximity map for non-dividing NB; Green: Jupiter::GFP. Note the bright flashes of red corresponding to NB divisions Scale bar 10 um.

**Supplemental Movie 3:** (See Figure 1D) Tracking of GMCs in a live explanted larval brain under extended time-lapse imaging conditions, collected at 2 minute intervals and displayed at 5 fps. Red: Histone::RFP; Green: Jupiter::GFP. Coloured dots: Tracked GMC candidates, using CytoCensus and trackpy, identified by colour. GMC divisions are visible (e.g. 21s top left, 1:04 centre). Scale bar 10 um.

