## Supplemental Material for "CytoCensus: mapping cell identity and division in tissues and organs using machine learning"

### SUPPLEMENTAL INFORMATION

#### Application of an ensemble of decision trees framework to identify and quantitate cell classes in 4D

With the development of CytoCensus, we aim to make identification of cell types in multi-dimensional image sets as straightforward as possible. We do so by asking the user to identify cell centres for a small number of 2D image slices. Using the cell centre annotation and the estimated size of the cells, we create an initial 'proximity map' of cell centres, similar in concept to density kernel estimation approaches (Waithe, *et al*., 2016, Fiaschi *et al*., 2012, Lempitsky & Zisserman, 2010). The advantage of using such methods to detect and count cells has been previously documented, but these methods have not been extended to the case of 3D cell centres with 2D annotations (Kainz *et al.* 2015, Waithe *et al.* 2016). Once we have the proximity maps of the cell centres, we then apply a series of image filters, which pull out image features such as edges, and try to use these features to predict a new proximity map of cell centres. We generate this new proximity map using a machine learning algorithm known as an "ensemble of decision trees" (Breiman, 2001; Breiman *et al*., 1984), which creates a series of "decision trees" that individually predict poorly, but averaged together are a strong predictor. Once we have the new proximity maps, the location of cell centres in 3D is inferred from the 2D predictions by applying a 3D Hessian filter (see later), which enhances the detections and resolves their coordinates in the additional dimension.

The CytoCensus software has three main components in its workflow: The 2D training and evaluation algorithm, the 3D object finding algorithm and the 3D ROI drawing and interpolation algorithms. The software is written in python and includes a Graphical User Interface (GUI) written using the PyQt library. The 2D training and evaluation algorithm utilises an ensemble of random decision trees and a bank of filters which utilise the matplotlib, scipy, scikit-learn and scikit-image libraries (Jones et al., 2001; Hunter J. D., 2007; Pedregosa F. et al. 2011; van der Walt S., 2014). For the 2D training and evaluation algorithm, the user must provide suitable images and make annotations indicating the locations of features, objects or cells of interest within defined regions. The user annotates 2-D sections of 3D image volumes and defines rectangular regions which encapsulate areas containing cells or features of interest or just background. There are N image volumes ($I_{i=1}$, $I_2$, $I_3$,…, $I_N$) in the training set and M annotation sections where M > 1 ($A_{j=1}$, $A_2$, $A_3$,…, $A_M$). Each annotation contains a region of interest ($R_{j=1}$, $R_2$, $R_3$,…, $R_M$) and also a set of corresponding points ($P_{j=1}$, $P_2$, $P_3$,…,$P_M$) with one or more dot/points Pj = {$pt_{c=1}$, $pt_2$,…,$pt_C$} or no points if the region only contains background. It is worth noting that providing sufficient area that does not contain cells is important for minimising false positives. As the model is designed to distinguish cells from the background it maybe appropriate to annotate regions as empty so as to acclimatise the model to the background. The points and regions are supplied by the user as they label the centroid locations of cells or objects within the image plane of interest. For each annotation we produce a centre-of-mass representation ($F_{j=1}$,$F_2$, $F_3$…,$F_M$) which for each pixel (p) is defined as the maximum value of all the Gaussian kernels (N) centred on dot annotations which overlap this pixel:

$$\forall p \in R_j,\ \ F_j^0(p) = max\left[\mathcal{N}(p; pt, \sigma^2\mathbf{1_{2x2}}), \forall pt \in \mathrm{P_j}\right]$$

and σ = [$\sigma_x$, $\sigma_y$]. The kernel is isotropic ($\sigma_x = \sigma_y$) as long the features or cells of interest are roughly spherical. For this application we recommend choosing a sigma which is smaller than the radius of the cells or features. The Gaussian will weight pixels in the centre of cells more highly than those towards the edges or in the background. Finding the maximum pixel, rather than summing pixels amongst all the overlapping Gaussians, ensures that pixels at the edges of objects, but overlapping, are not more highly weighted than pixels that are central and represent the centre of the cells, allowing better separation of close objects.

For each pixel in the annotation region we calculate a feature vector which describes the corresponding image pixels. Each descriptor of the feature vector is created through processing of the input image or volume with one of a bank of filters which includes: Gaussian, Gaussian Gradient Magnitude, Laplacian of Gaussian, and the minimum and maximum eigenvalues of curvature (Fiaschi *et al.*, 2012). These filter kernels are applied at multiple scales (sigma = 1,2,4,8,16) to aggregate data from the surrounding pixels into the feature descriptor at that specific pixel. This scale range was appropriate for all the cases used in this study and were not changed.

Once training data has been supplied by the user and the pixel features calculated, an ensemble of random decision trees is used to learn the association between input pixels and the “proximity map” centre-of-mass representation (Geurts *et al.*, 2006). The decision tree framework was parameterised as follows: the data was sampled at a rate of 1/5 from the input regions, with 30 trees generated during training, with a depth of 10 levels and a minimum split condition of 20 samples for each node. At each node n/3 features were considered. Once trained, the decision tree framework can be applied to unseen images (without user annotation), requiring only input features to be calculated. Evaluation of images produces a centre-of-mass representation of where the cell centres are located, highly similar to the representation used during training.

The 3D object finding algorithm is applied to the output images of the random decision tree framework and involves multiple steps. Firstly the output images of the decision framework are rearranged into a 3D volume, this provides a representation of the proximity of cell centres in 3D. To facilitate the object identification we next apply a determinant of Hessian blob detector which smooths our signal and also enhances objects of a specific size (Lindeberg, 1994). Using this filter greatly simplifies our cell identification procedure although some idea of the size of the object is required, $h = [h_x, h_y, h_z]$ (where $h_x = h_y$ if the object is spherical in two dimensions and $h_x = h_y = h_z$ if the object is spherical in three dimensions). Finally a 3D maxima finding algorithm is used to identify the centroid locations of the enhanced objects present in the Hessian filtered image (Gao and Kilfoil, 2009). A simple threshold is used to set the sensitivity to detected maxima.

To allow for selective application of the 3D counting algorithm in distinct regions of a tissue (for instance the primitive streak in Figure 6), and over time, a novel Region Of Interest (ROI) interpolation algorithm was introduced. The user defines a ROI by clicking points around an area of interest in a single image (e.g. top of tissue region). The user then defines another region either at the other end of the object (e.g. bottom of tissue region) or partially through the region. The algorithm can then interpolate between these user-defined ROI to create a ROI for each frame in the image-volume. The User can then repeat this process in subsequent time-frames, and the algorithm will interpolate the ROI between frames creating a smooth transition which can be tweaked through the addition of further user defined regions to smoothly follow a 3D region of the tissue over time. The interpolation is performed using bilinear interpolation of points sampled uniformly along the user defined ROI. Objects or cells with a centroid position within the tissue region can then be filtered from the image volume allowing for selective counting and location of cells over-time within the defined region.

#### Algorithm Validation and Comparison Datasets

Validation of algorithm performance is critical in developing an effective tool. We used both real and artificial data sets to assess performance. For our baseline performance tests of CytoCensus, we quantified the number and location of NBs identified in five time-points from a movie sequence. We then compared the output from CytoCensus to that of other algorithms applied to the same test dataset. In each case, we attempted to optimise the parameters used, based, whenever possible, on the published information on two time-points that were not used for final evaluation. For TrackMate detections we used detection diameter of 20 pixels, and 5 conservatively set filters (standard deviation, max intensity, mean intensity, contrast). For RACE we used the histone (nuclear) marker for the seeds, and set parameters at default except for (Max segmentation area 100, min 3D area 20, Closing 2-6, Threshold 0.0002, H-maxima 30). For Ilastik, we used features from (sigma 0.3, 1.0, 3.5, 10) for color, edge and intensity. For FIJI WEKA, we used default parameters, followed by filtering to remove small objects. For CytoCensus we used default parameters, except for object size, which was set at radius 8.

To carry out a direct comparison of algorithm performance we used artificial "neutral challenge" datasets of highly clustered (75%) synthetic cells, in 3D, with a low signal to noise ratio (SNR), obtained from the Broad Bioimage Benchmark Collection (image set BBBC024vl: Svoboda et al., 2009). This data has an absolute ground truth and provides a good measure of comparison for the performance of different algorithms. As CytoCensus is designed to identify cell centres, the ground truth for the Neutral Challenge data and Ilastik segmentation results were adapted to report estimated cell centres (centroids) rather than segmented boundaries, before carrying out comparison of algorithm performance. In both cases, Ilastik and CytoCensus were trained on a single image, parameters optimised over 5 image, and performance evaluated over the remaining (25) images. For the neuroblast dataset, detections within 1/2 a neuroblast radius were considered correct. For the BBBC dataset, detections were considered correct if they were within a stricter 4 pixel radius (~1/8 cell size). At the more generous 1/2 a cell size, CytoCensus reached perfect precision and recall, but Ilastik's F1-score remained below 0.4 primarily due to the problem of merged cells.

##### F1-score: Objective Analysis of Algorithm Performance

To quantitate how CytoCensus analysis of complex multidimensional image data is performing and to compare performance to other freely available programs, we made use of the weighted mean of the true and false positive identification rates, known as the F1-score (maximum value 1.0; Chinchor, 1992) (Figures 3 and S4, Table 1). F1-score is intuitively similar to accuracy: strictly it is the harmonic mean of the fraction of detections that were correction (precision) and the fraction of cells correctly identified (recall). This metric, therefore, takes into account both false positives and false negatives. Such analysis is particularly useful in parameter determination for optimum algorithm performance in applying to experimental data sets and in optimising algorithms for systematic comparison on test data.

**Parameter Optimisation for Best Performance**

The algorithm underlying CytoCensus requires a range of parameters to be set, described in detail above. To simplify usage, we set most most parameters to reasonable default values. Parameters, such as the threshold setting, were assessed systematically, using the objective measures of performance described above with various data sets. This approach helped us to define which parameters should be fixed and which need to be user-modified. Details of user defined parameters and how to assess and set appropriate values are documented in the User Manual. The value of Sigma, which sets the scale of the object of interest, is particularly critical for detection. Optimum Sigma value was assessed systematically and the optimum found to be slightly smaller than the size of the cell type of interest (Figure S4A”), this parameter was subsequently defined as “Object Size” (in pixels).

The level of training required is also important in supervised machine-learning approaches. We assessed the level of training required to achieve good detection of cell types of interest with different datasets (Figure S4A’). In all cases tested, we found that successful identification of NBs or progeny required minimal user training (of the order of tens of examples on only a few image planes) and increasing training gave only marginal improvement (Figure S4A’). This is advantageous for quickly annotating datasets, but may limit the flexibility for learning in particularly difficult and complex cases.

**Cell identification**

To facilitate development of CytoCensus for the study of the larval brain, we initially tested performance on multichannel 3D image datasets of fixed material where NBs and GMC's were defined by specific immunostaining (for Ase and Dpn; Neumüller *et al.,* 2011; Bayraktar *et al*., 2010; Boone & Doe, 2008) (Figure S3A, B). We confirmed that, given these ideal markers, NB's and GMC's could be identified. After this initial development we extended the application of CytoCensus to our live cell imaging data with generic cytological markers. To show that cell types could be recognised correctly with the generic marker combination used for live imaging, we carried out specific immunostaining on Jupiter::GFP / Histone::RFP expressing larval brains. Manual and automated annotations, first based upon generic labels alone, were scored against identification using the specific labelling (Figure S3C). The results of this assessment show that, for NB (96% ± 4 Dpn positive, n = 12, 3 repeats) and their progeny (92% ± 2 Pros positive, n = 189, 3 repeats), our imaging of the generic labels supports identification of NB and progeny by CytoCensus after training (Figure S3D). Using a combination of these different datasets we refined the workflow of CytoCensus and optimised the key parameters of the algorithm.

**Image pre-processing and Downstream Data Analysis**

CytoCensus outputs proximity map images and object centre XYZ co-ordinates, both of which may be used as the starting points for further data analysis pipelines, for example as seeds in watershed segmentation to determine cell areas, volumes or quantitate fluorescence intensity. CytoCensus outputs (tif files) which can easily be passed to Fiji (ImageJ) or custom analysis scripts. In this study we illustrate several examples of extending data analysis using outputs from CytoCensus.

In our analysis of NB division rates, we generate plots of cell division over time for individual NB from the map output and NB centre coordinates (Figures 5A,B and S6). Here we used a custom trackpy based python script to track individual NBs over time, however, similar results can be achieved simply, using ImageJ ROI tools, or at scale using TrackMate (Figure S6A). For each of these tracks we follow the changes in the dividing NB proximity map that correspond to division.Robustness of the division plots is further improved by subtracting a moving average over about 20 image frames, which removes spatial differences in the background of the probability density maps. For analysis of long imaging series, such as multiple NB divisions, it is important to follow individual cells over time. For such analysis, it is necessary to spatially register the individual Z stacks across time, to correct for image drift due to movements in culture, prior to applying CytoCensus. This was achieved with an ITK based python script (http://www.simpleitk.org/

SimpleITK/resources/software.html), but similar results can be achieved using the Correct 3D drift plugin in ImageJ (it is crucial to use a high noise threshold, ignoring low value pixels, as this approach is sensitive to noise). Similar approaches were employed in our analysis of GMC cell cycle length (Figure 4D). In the challenging case of GMCs it was necessary to explicitly track cells as they moved significantly during development of the brain. To achieve this, estimated GMC centres were passed to a trackpy (Allan et al. 2016), based custom python script to perform linkage analysis and track each individual GMC over time. Similar results may be achieved using TrackMate in FIJI (Tinevez *et al*., 2016). Python scripts for further analysis are available on the CytoCensus Github page.

### SUPPLEMENTAL FIGURES

**Fig.S1:** *Ex-vivo* larval brains continue to develop in culture

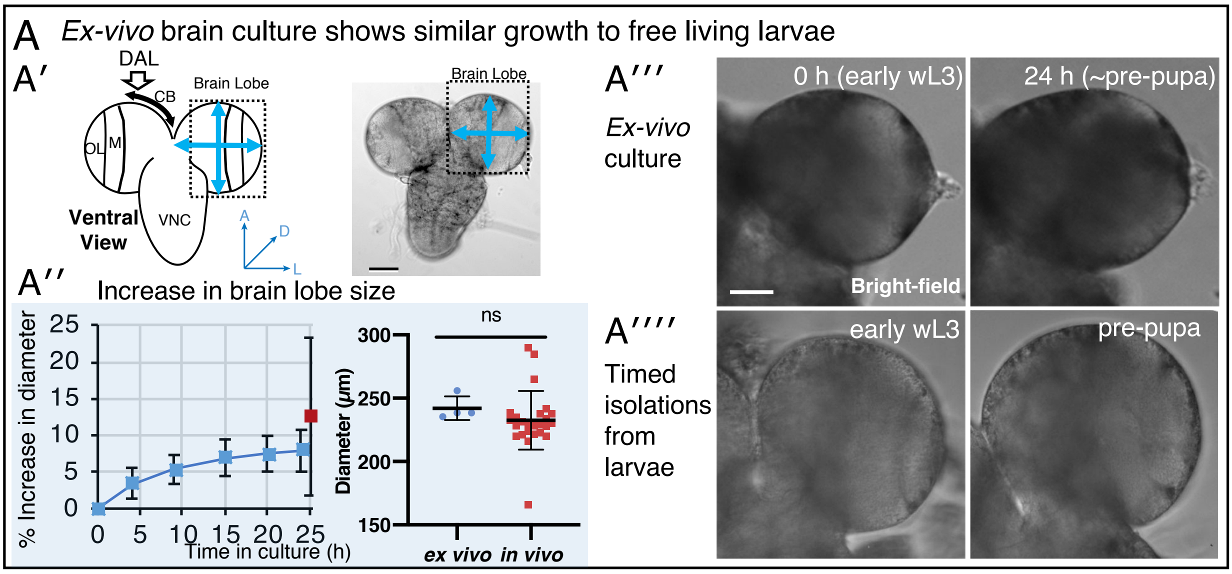

**Figure S1. *Ex-vivo* larval brains continue to develop in culture. Related to Figure 1 A′**) Left panel: diagrammatic overview of the *Drosophila* wL3 brain highlighting the dorso-anterior-lateral region of the central brain which contains neuronal stem cells of the type I NB lineage. CB central brain; OL optic lobe; M medulla; VNC ventral nerve chord; DAL dorsal anterior lateral region. Right panel: an *ex-vivo* brain imaged in bright field, the blue arrows indicate measurements taken of brain lobe diameter to assess growth in culture. **A′′**) Analysis of the increase in brain lobe size in culture compared to *in-vivo*. The left plot shows *ex-vivo* cultured brain lobe diameter increase for Jupiter::GFP; Histone::RFP L3 brains over 24 h (blue trace, n = 3) under wide-field fluorescence imaging conditions. The single red datapoint shows average lobe diameter (n = 13) for freshly isolated brains from free living larvae at the end of wL3, corresponding to the stage expected for 24 h culture. The right hand plot directly compares brain lobe diameter for 24 h cultured and freshly isolated brains from free living larvae at the end of wL3 (ns, Mann-Whitney test, *ex-vivo* n = 4; *in-vivo* n = 25). **A′′′)** and **A′′′′)**, bright field images at two time-points during culture and examples of freshly dissected free living larvae at corresponding developmental stages, respectively. Scale bars 50 $\mu$m;

**Fig. S2**: CytoCensus graphical user interface

A User interface snapshots

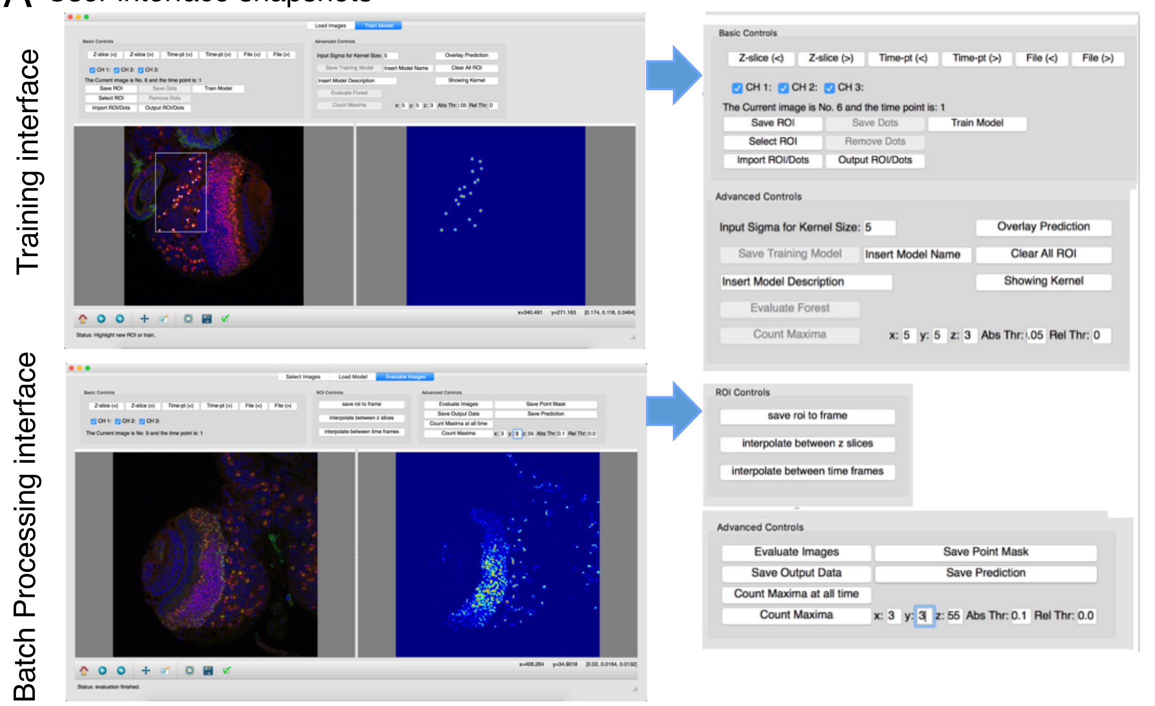


B Program output data

| | Filename: | Time point: | Predicted count: |
|---|---|---|---|
| Thu Jun 2 17:17:47 2016 | 20150206_DAPI_Dpn568_Ase647_60x_21Stacks_WT03_BRG.oib.tif | 1 | 330 |

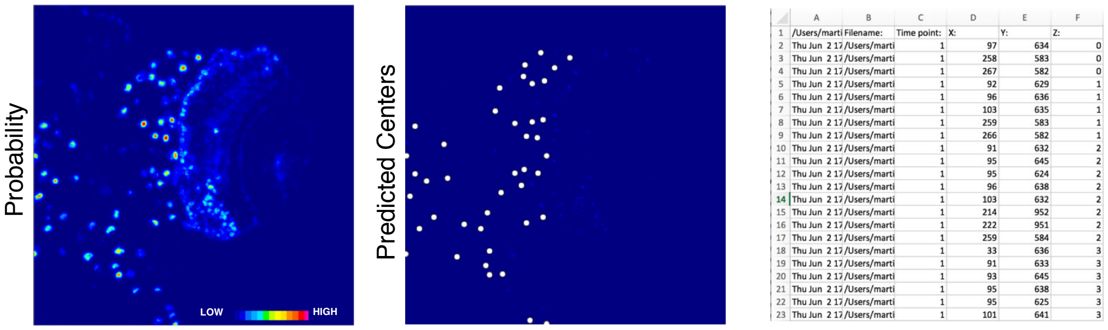


**Figure S2. CytoCensus graphical user interface. Related to Figure 2 A**) Snapshot of the user interface for 'Training '(above) and 'Evaluate' batch processing (below), with the user defined settings enlarged to the right (for more details see the '**User Manual**' associated with the Supplemental material). (**B**) Output results summary for automated detection of a cell class: count; probability map, predicted centres overlay; XY co-ordinates for predicted centres.

**Fig. S3**: Validation of cell identification by CytoCensus

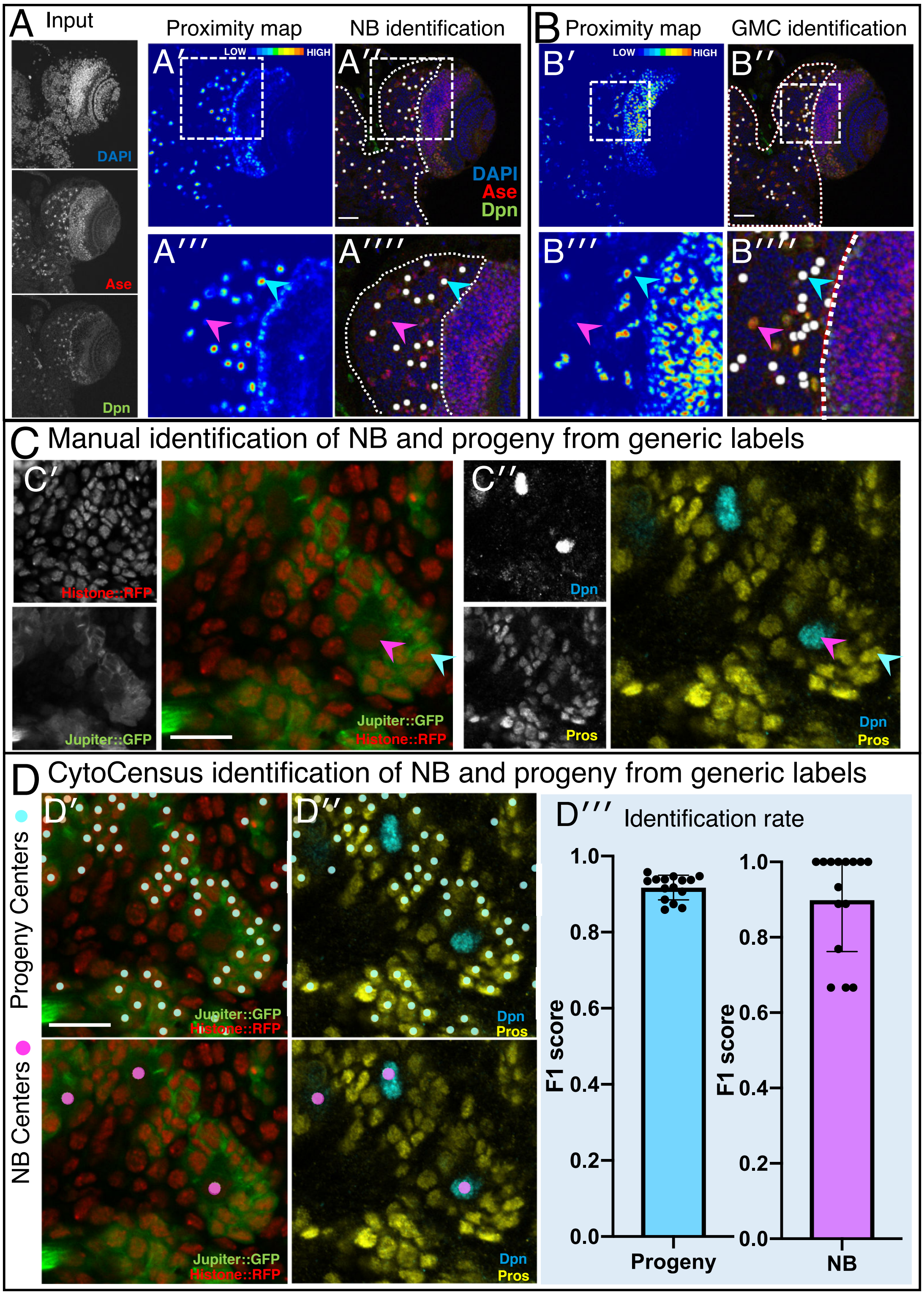

**Figure S3. CytoCensus identification of cell types. Related to Figure 3.** Examples of automated detection of **A**) Type-I NBs and **B**) GMCs, respectively, from a 3D multichannel image set for a fixed WT larval brain labelled with DAPI, anti-Ase and anti-Dpn. In both cases training was performed by single click annotation within a user defined region of interest (ROI, dashed lines in **A″** and **B″**) to identify the cell class of interest. The resultant “proximity” maps (**A′** and **B′**) and corresponding cell class identifications (e.g. **A″** and **B″**)were evaluated manually to assess the success of training. The zoomed regions (corresponding to the dashed white boxes) show examples of correct identification (cyan arrowheads) and false negative (magenta arrowheads) of NB in **A′ᵛ**) and GMC's in **B′ᵛ**). **C**) Manual identification of cell classes from generic labels: fixed Jupiter::GFP, Histone::RFP labelled brain (**C′**), immuno-labelled for Dpn and Pros (**C′′**) to permit unequivocal identification of NB and their progeny. **D**) Validation of CytoCensus identification of NB and progeny from generic cytological makers. **D′**) Cell centre predictions are shown from CytoCensus analysis of the dataset from (**C′**) with generic markers. **D′′**) Corresponding identifications of NB and progeny based upon Dpn and Pros markers. **D′′′**) Plot showing that identification based upon Jupiter::GFP, Histone::RFP labelling alone effectively identifies NB and progeny compared to identification from Dpn and Pros labels: 96% ± 4 NB identification (n=12, 3 repeats) and 92% ± 2 progeny identification (n=189, 3 repeats). Scale bars **A″** and **B″** 50 μm; **C′** and **D′** 20 $\mu$m.

**Fig. S4**: Optimisation of CytoCensus parameters for cell center identification

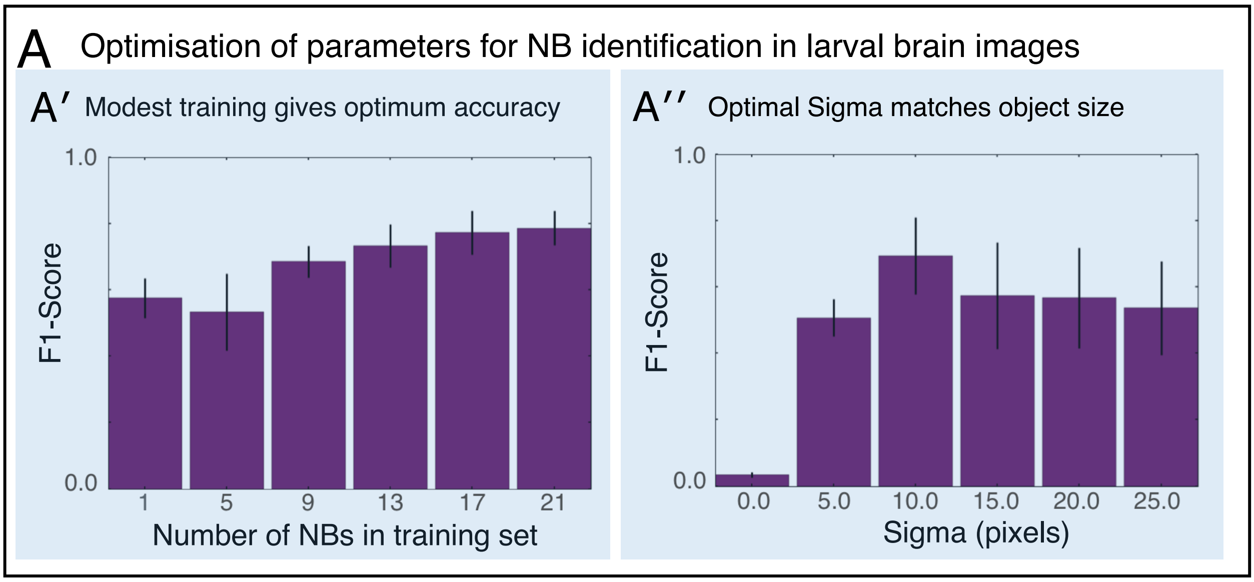

**Figure S4. Optimisation of CytoCensus parameters for cell centre identification. Related to Figure 3. A'** The F1-score (see Supplemental Information) was plotted for different levels of training from annotation of a single NB cell, up to annotations of 21 individual NB. As training examples are added, results improve but rapidly approach maximum with little training **A"** F1-score for varying sizes of sigma (radius of cell detection in pixels). Performance is optimal at sigma = 10, which is slightly smaller than the radius of a NB.

**Fig. S5**: Loss of Syp causes enlarged larval brains

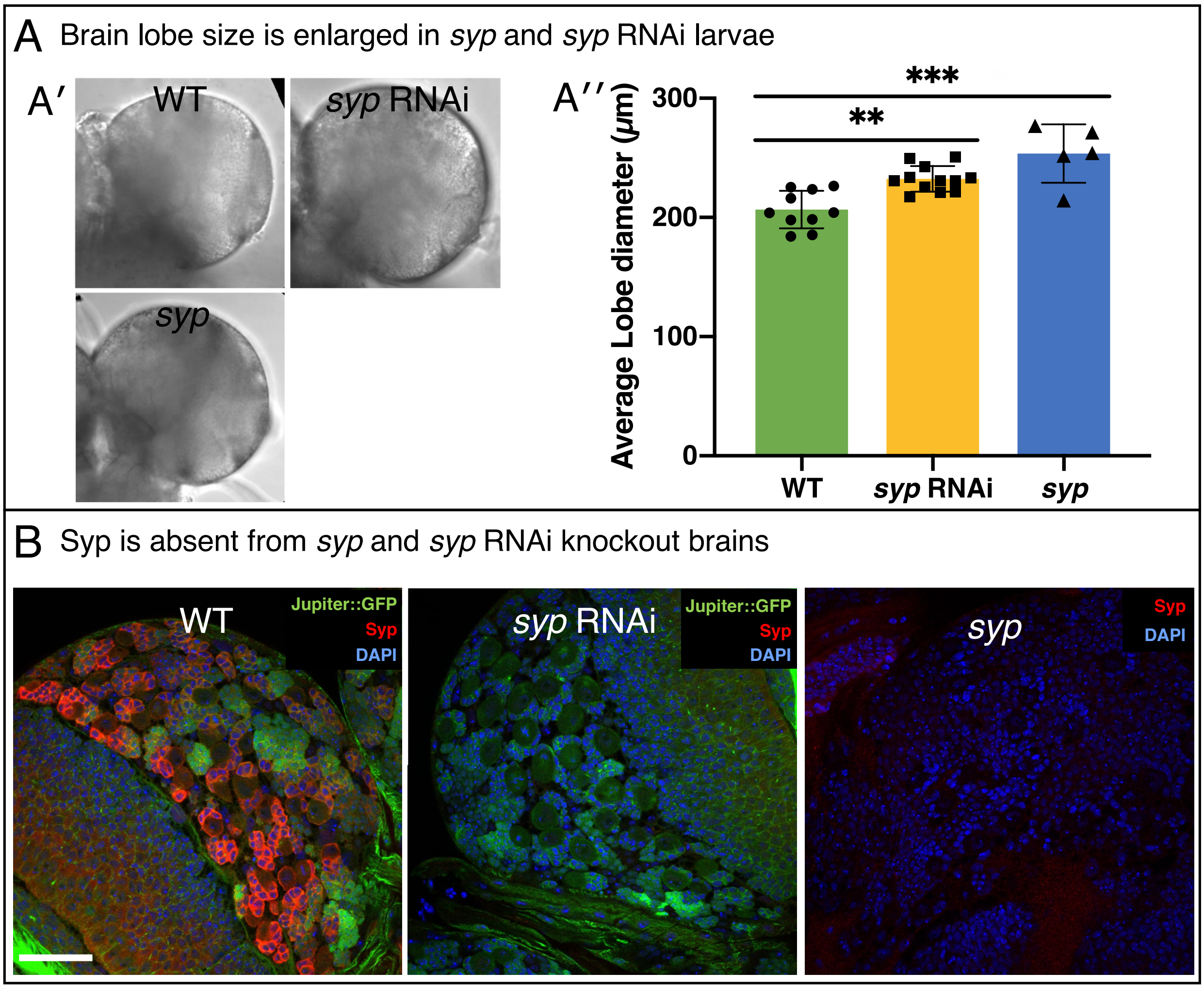

**Figure S5. Loss of Syp causes enlarged larval brains. Related to Figure 4. A**) Comparison of brain lobe diameter **A')** Brightfield images of WT, *syp* and *syp* RNAi **A'')** Average of 2 manual measurements/lobe (as per Figure S1A). Lobe diameter is increased in *syp* (WT vs *syp,* $p<0.0001$, t-test with Tukey correction, n=12), and *syp* RNAi (WT vs *syp* RNAi p=0.0022, t-test with Tukey correction, n=5) **B**) Immunofluorescence analysis shows absence of Syp expression in *syp* RNAi, and *syp* mutant brains. Brains are labelled with DAPI, anti-Syp and Jupiter::GFP. Scale bar 25 μm.

**Fig. S6**: Following NB divisions with CytoCensus

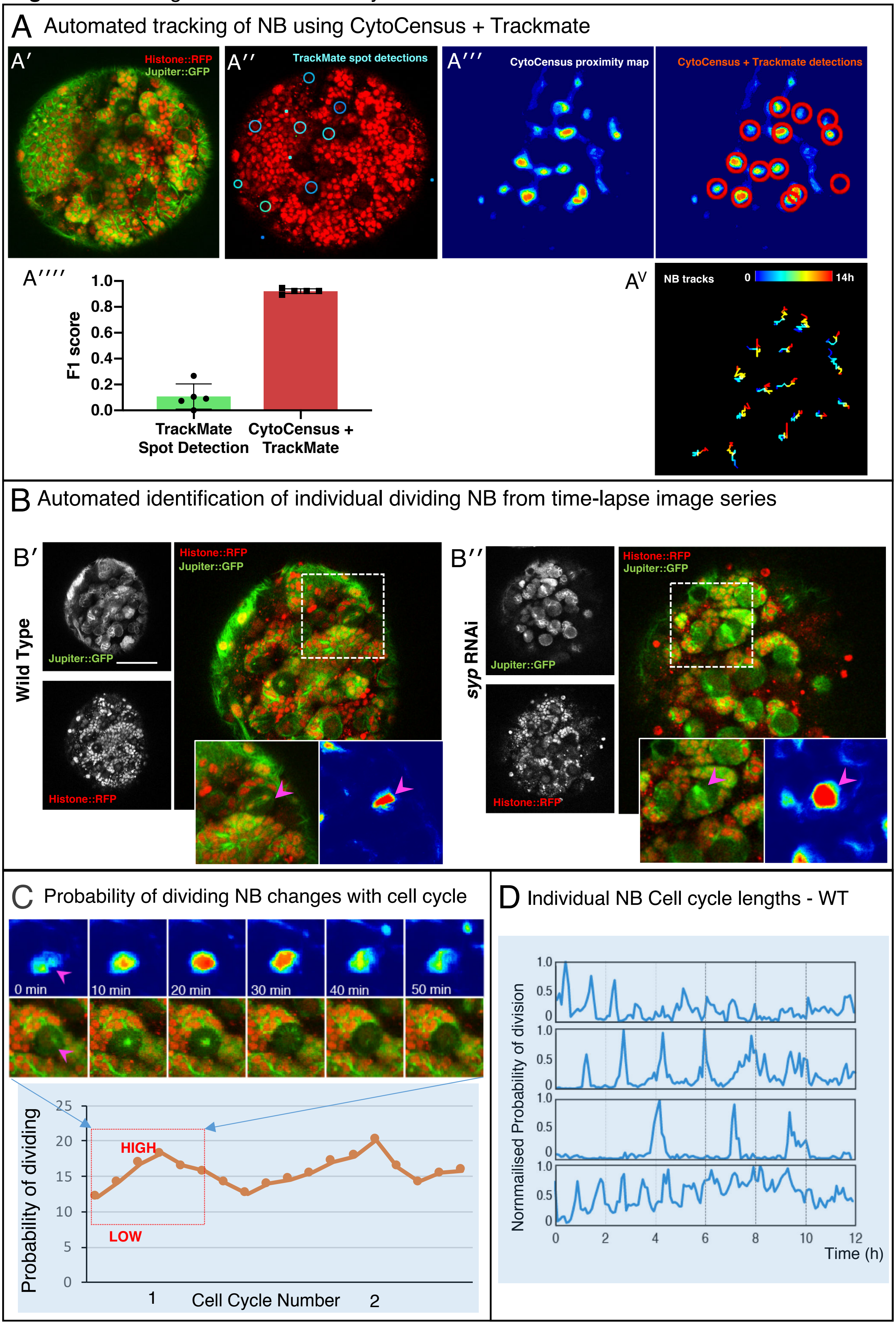

**Figure S6 Following NB division with CytoCensus. Related to Figure 5. A)** Tracking of NB with TrackMate and CytoCensus **A′-A″)** Raw image, and histone-based detections of NB (cyan circles) from TrackMate **A‴)** CytoCensus NB proximity map, and corresponding TrackMate detections (red circles) **A⁗**) Graph of F1-score of raw TrackMate detections and CytoCensus proximity map with TrackMate detections **$A^{V}$)** NB tracks that span the whole 14h movie from CytoCensus + TrackMate **B**) Automated identification of individual dividing NB for time-lapse series using the probability density map output of CytoCensus. **B′-B″**) Show WT and *syp* RNAi brains, with inset highlighting an individual dividing NB (marked with anti-Ase, anti-Dpn and DAPI) and their corresponding proximity maps. Scale bar is 50 µm. **C**) Change in proximity score (probability of division) plotted over time for an individual WT NB undergoing division: upper panels are the confocal images and corresponding proximity maps; lower panel is a plot of probability covering two cell cycles, the region corresponding to the images above is highlighted. **D)** Series of plots showing different NB over time, and the changes in the probability of dividing (B) as they progress through the cell cycle.

**Fig. S7**: CytoCensus detections of PGCs in Mouse Embryos

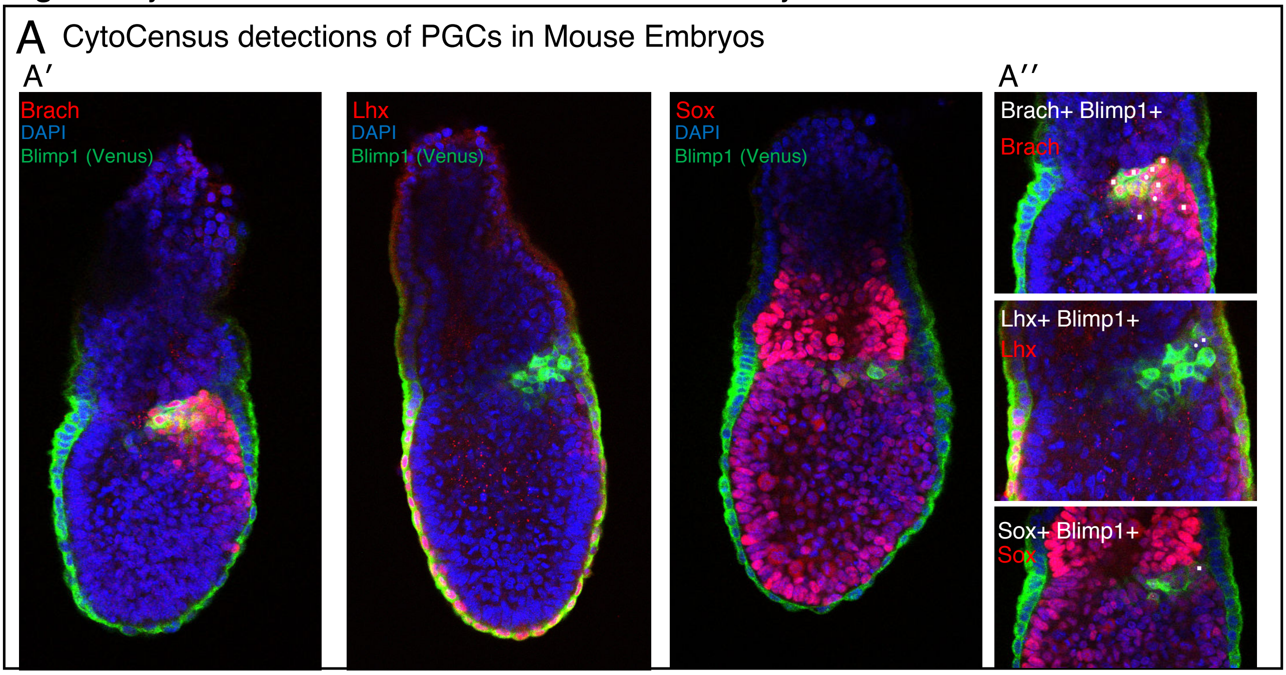


**Figure S7 CytoCensus detections of cells in Mouse embryos Related to Figure 6. A′)** Image plane through centre of each embryo used for quantification illustrating the different distributions of TF⁺ cells. Images marker with DAPI, Blimp1-Venus and antibody against (Brach, Lhx, Sox) **A″)** TF⁺ (Brach, Lhx, Sox) PGC detections in the proximal posterior epiblast (PPE), quantified in Figure 6B

### User Manual: CytoCensus Quickstart Guide

CytoCensus is intended to allow biologists to identify and count cells and objects of interest in complex 3D microscopy images, without detailed knowledge of image analysis. Specifically, the program is targeted at users with large numbers (>10) of microscopy images or large time-lapse datasets, and asks the user to do some very simple annotations (training the program) to identify the cells of interest. Software downloads and test data sets may be found at: https://github.com/hailstonem/CytoCensus/releases/.

#### What can I do with CytoCensus?

- **Count cells/objects in 3D** – if they are approximately round.
- **Find cells/objects** of a particular diameter.
- **Identify cell/object centre**s i.e. XYZ coordinates.
- **Determine numbers and locations over time-lapse** image series.
- Determine object counts within a **user defined regions of interest (ROI)** over time.
- Use output values to compare relative numbers for different object classes (or subclasses).

With a little FIJI/ImageJ knowledge, CytoCensus outputs can be used to:

- **Determine average intensity** across a class of cells.
- **Track cells** over time, using CytoCensus probability maps as input to the ImageJ (FIJI) TrackMate plugin, to track previously untrackable cell classes.

#### What can't I do with CytoCensus?

- **Identify things than you cannot recognise.** If you cannot find the objects/cells of interest to train the program, CytoCensus will not work.
- **Find objects of extremely variable size.** CytoCensus allows some variation in size, but within limits - if you have a large set of cells and a small set, you should train separate models to recognise each.

- **Trace long, thin, or branching cells, such as neurons.** However, you can use CytoCensus to identify roughly round features of such cells, such as cell bodies or nuclei, and pass those co-ordinates to other analysis tools.
- **Measure cell areas\volumes directly.** Unlike segmentation tools, such as Ilastik, CytoCensus determines cell centres, not their edges (boundaries). To measure cell areas/ volumes it is possible to use the XYZ centres output from CytoCensus as the starting point in an image analysis pipeline.
- **Measure total fluorescence of cells directly.** As CytoCensus does not identify boundaries, it cannot determine the extent of the cell, therefore, total fluorescence cannot be measured directly. However, as described for cell volume above, CytoCensus outputs may be used as the starting point for an image analysis pipeline.

#### What kind of data should I use?

CytoCensus is designed to work with multiple 3D image files, with multiple z-slices, multiple channels, and/or multiple timepoints. CytoCensus requires TIFF files. If your data is in another format, convert it using the excellent FIJI/bioformats plugin found at https://imagej.net/Fiji/Downloads (Linkert *et al.*, 2010).

#### How do I use CytoCensus?

CytoCensus is best used on a computer with 16GB or more RAM with a mouse. CytoCensus works with a PC, Mac or Linux platform.

#### SETUP

1. Download the latest version of CytoCensus from https://github.com/hailstonem/CytoCensus/releases/. You will need both the 'CytoCensus_train.app' and 'Cytocensus_evaluate.app'.
2. Run 'CytoCensus_Train'.

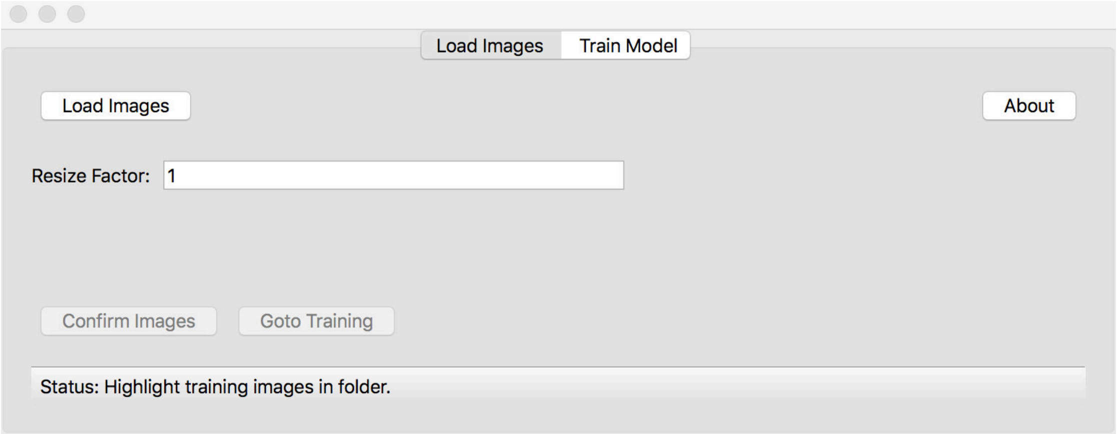


3. **Load images** (TIF/TIFF files supported only) - 3D images should be stacks or hyperstacks: https://imagej.nih.gov/ij/docs/guide/146-8.html
    a. If your images are in another format, use ImageJ/FIJI to convert them https://imagej.net/Fiji/Downloads
    b. You may choose multiple images files but at this point it is unnecessary to load your whole image dataset, simply load enough to represent any variation in your images.
4. If your images are large (>512x512pixels and/or z>50), and the objects you want to look at are also large (>20 pixels), choose a '**Resize factor**' (e.g. 2,4,5) before you load an image. This will speed up loading and processing, but setting this too high will reduce your image quality.
5. Set '**Sampling**': 5 is the default and generally works. Decreasing this to 2 (minimum = 1) can provide better accuracy with rare and small objects (<5pixels). 'Sampling' sets how many pixels are used in calculations to determine the probability density values.

6. ‘**Feature set**’. Leave this at ‘Pyramid’ for most cases. Basic is faster, but requires you to set the ‘feature size’ parameter (see later). ‘Histogram equalised’ is slower, but can help if you image contrast varies a lot, for example over a time-course dataset.
7. ‘**Confirm Images**’ and ‘**Goto Training**’.

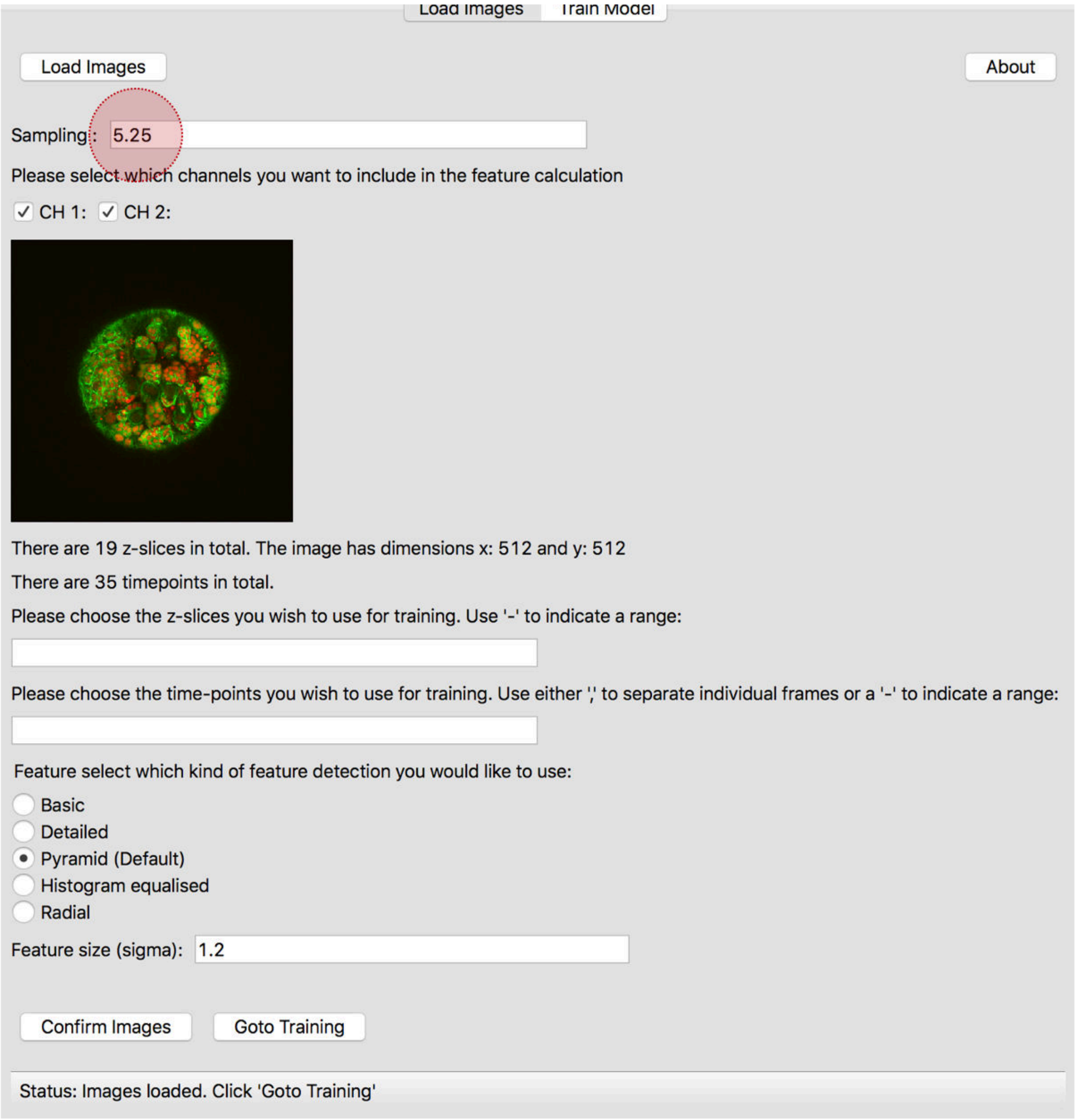

#### TRAINING

8. **Navigate** through your images using up down keys (z), left right keys (t), < > keys (files) or the corresponding buttons on the GUI.

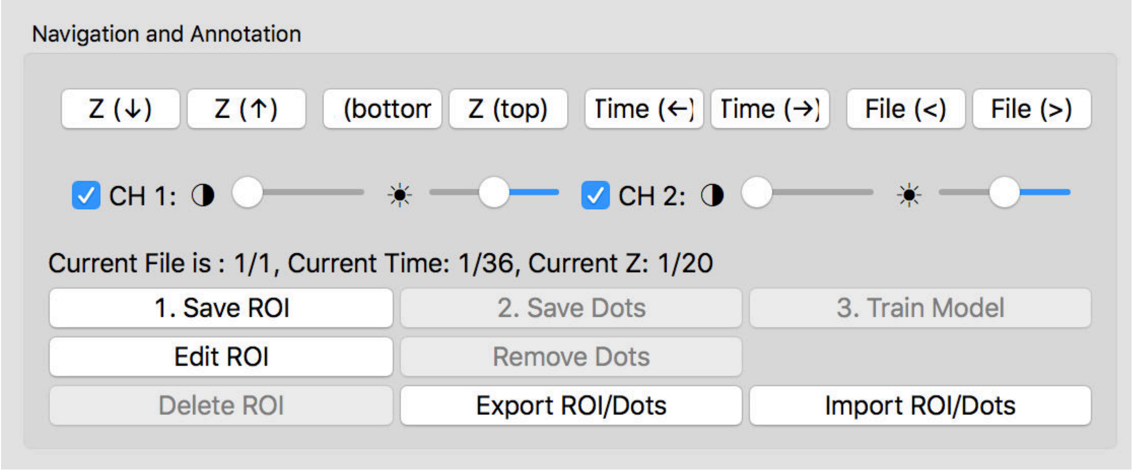


9. Find a z-slice with an example of your cell or object of interest. Adjust brightness / contrast using the 'sliders' if necessary.

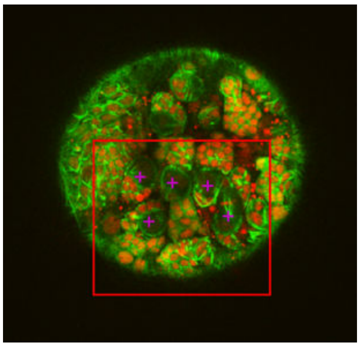

10. **Right-click and drag to make an ROI** containing a number of objects of interest and some background (parts of the image that are not the features of interest).
11. When happy with your ROI, **left-click your objects of interest** in it. If you are not happy, simply right-click and drag to redraw a new ROI. Note, boxes cannot be moved or resized. Don't change image/z or you'll lose your ROI. You can remove erroneous marked objects with the 'remove dots' command and then selecting the individual marks to remove.
12. **Save** your ROI: select '**1. Save ROI**'. Save dots: select '**2. Save Dots**'.

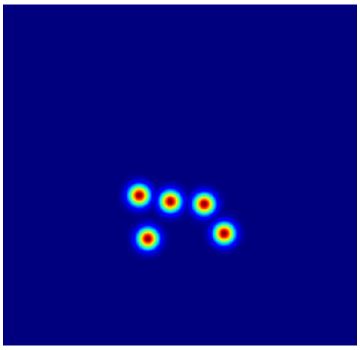

13. You should now see some points in the right hand plot. These should be roughly the same size as your object of interest. If they're much smaller, or barely visible, increase the '**Object Size'** parameter until the size roughly matches that of your objects. Do not worry too much if this isn't perfect, it can be changed later.

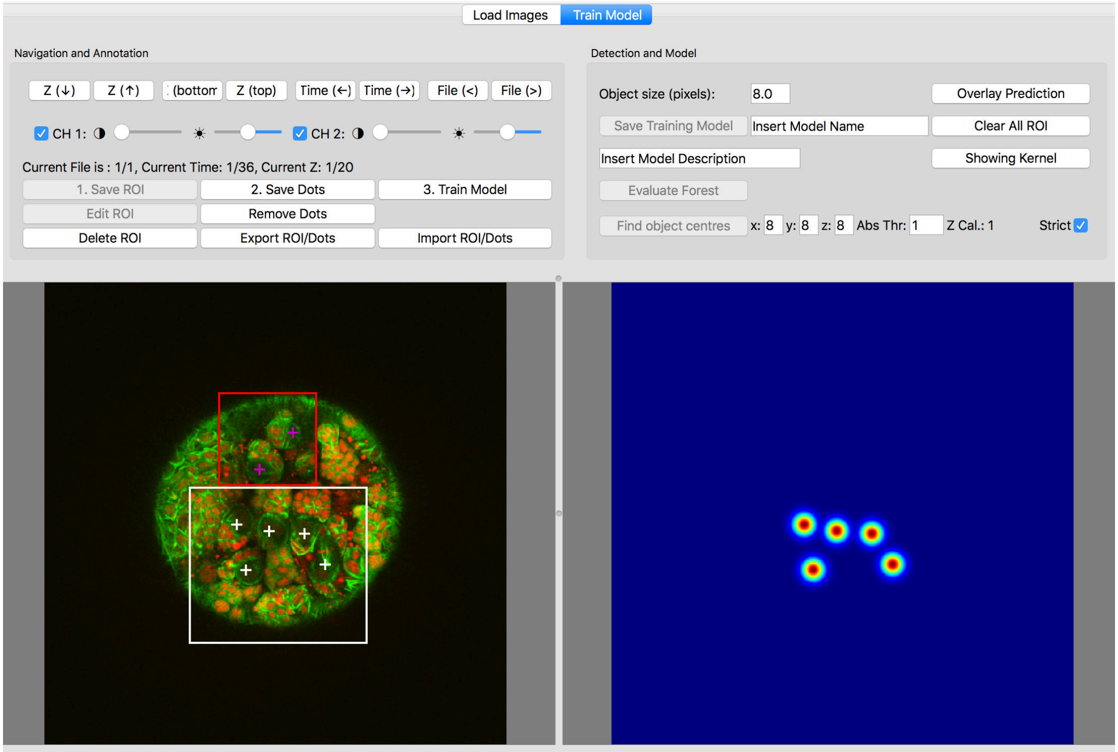


14. Add more training if desired (8-12), ideally in a different z/t. Note: adding multiple annotations in Z planes rather than timepoints is more efficient for the program but that results will only be displayed for the current Z-stack (all your training is used!).
15. When happy with your training select ‘**3. Train Model’.**

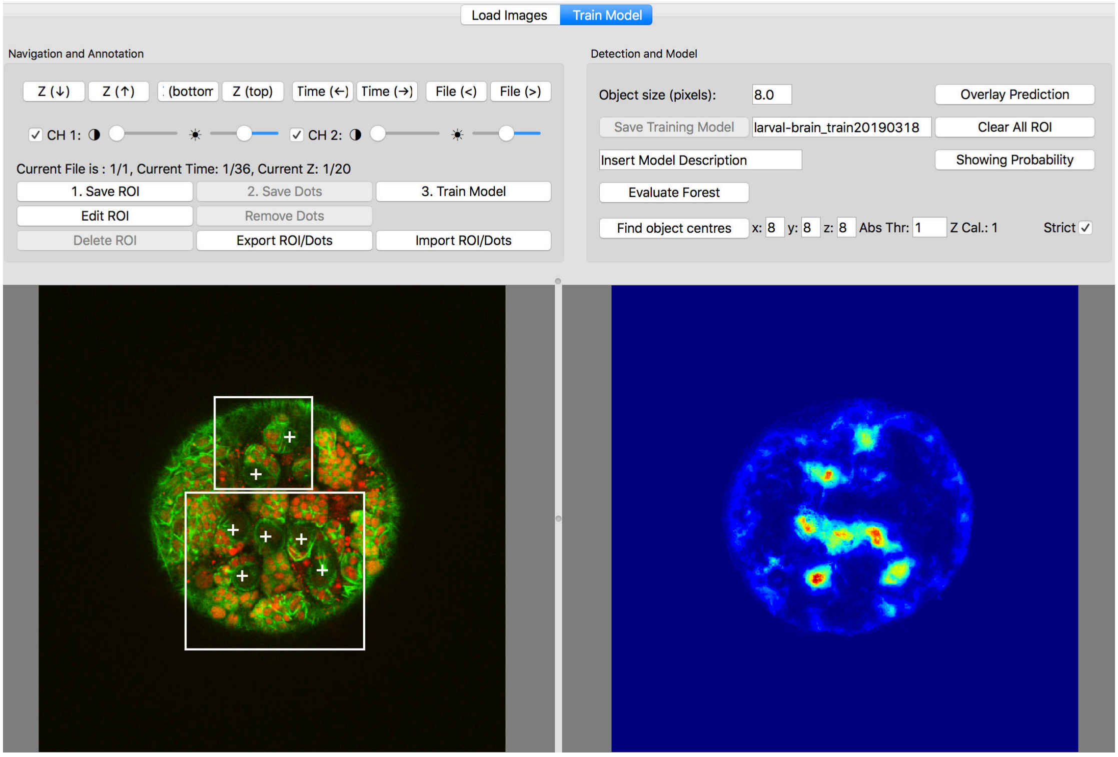

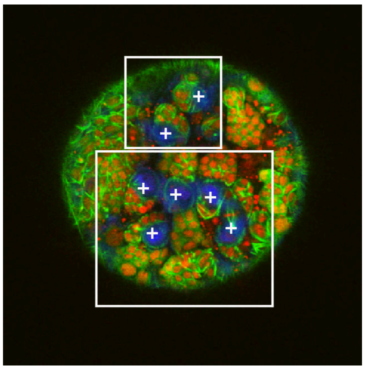

16. Use the **probability map** on the right to evaluate the training in the current z stack, you can also use the '**Overlay Prediction'** button to help relate the map to features (left panel). To examine the result in a different z-stack, use the '**Evaluate Forest'** button. In an ideal map there should be clear discrete regions of green/yellow/red corresponding to your objects of interest, and faint blue or very little elsewhere. Add more training as described in 8-14 to improve your training. Note that you can add more points, but also that you can make new ROIs with no features of interest in them if there is some background that is causing an issue.
17. When you are happy with the **probability map,** should progress to '**Find Object centres**'. This will attempt to find the highest intensity points of interest in 3D and is the hardest part to get right being quite variable depending on how consistent your objects are. There are **four parameters** that are important for this:

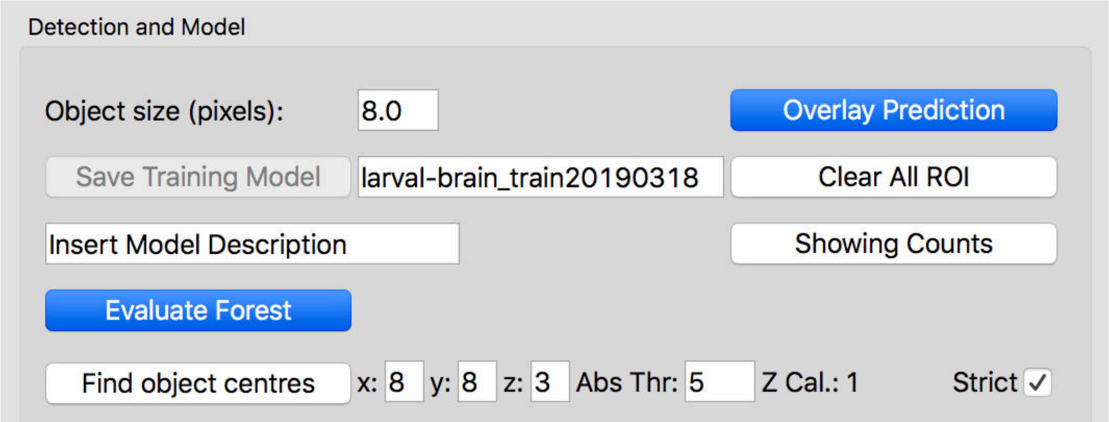


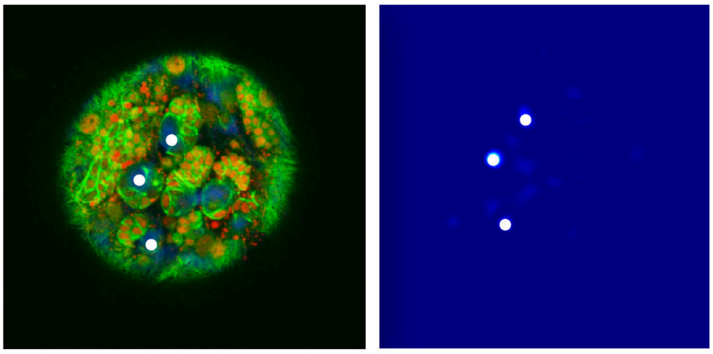

The thresholding, '**Abs Thr**', parameter is the most important: typically a value between 0 and 20 is best. The **x and y sizes** should be a good estimate based on the size of objects chosen earlier, but the **z size** can vary a lot. If the **z calibration** on the right is accurate (this is the difference between your x/y resolution and your z resolution) then matching it to the object size make sense. If you have 1um pixel size in x/y and take 5um steps, then z calibration would be 5. If your object size is 10 pixels, then, a reasonable value for x,y,z sizes is 10, but you may need to try changing the z size between 5 and 15 to see what gives the best result.

18. When you're happy with this, **choose a name** for your "Training Model" and the **Save Training Model.**

#### BATCH MODE (Evaluation of new data)

After training by the user, CytoCensus employs a collection of filters and scores and combines them to find features in the image that identify the user-defined cell centres. In this way, a series of transformations of the image data (referred to as the "**trained model**") are learned. The trained model is subsequently applied pixel by pixel to new data sets, outputting an estimated probability map of the cells of interest with their predicted cell centre co-ordinates.

19. The hard part is now done. Now open 'Cytocensus_evaluate.app'.
20. Load in your images as before (**Add images**). This time load the whole image dataset (i.e. leave the options for Z and T blank).

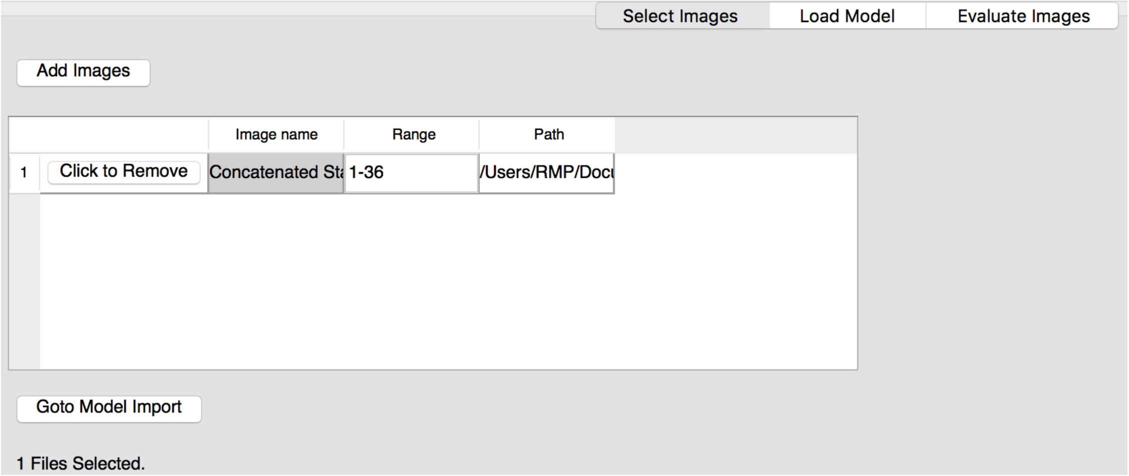


21. **'Goto Model Import**'. Note, take care in applying models across data sets, particularly, for example wild type (WT) vs mutant data sets. If there is a significant difference in data quality of the object (cell type) of interest between datasets, then retraining is required. However, if may be of value to apply training from a WT case to a mutant - if the cells of interest are not detected then there is likely a significant difference from WT - i.e. WT cells are not present in the mutant.

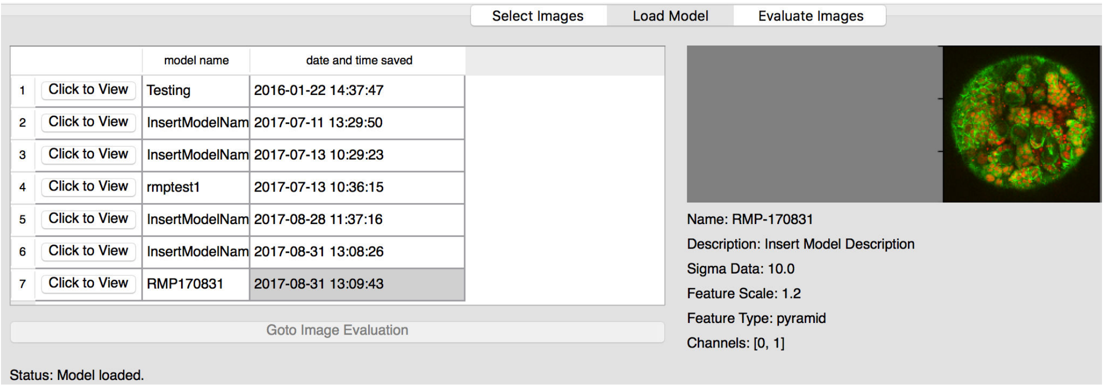


22. Click on your **saved Model and click to view it, then select 'load model'.**
23. Go to **'Image Evaluation'.**

**24. Evaluate Images.** This is the most time-consuming step, but does not require further user input. Our our data sets, evaluation takes a little longer than 5 minutes per 50 z-planes, so can be a couple of hours for a 150 time point movie, but this will depend on the computer you're running it on.

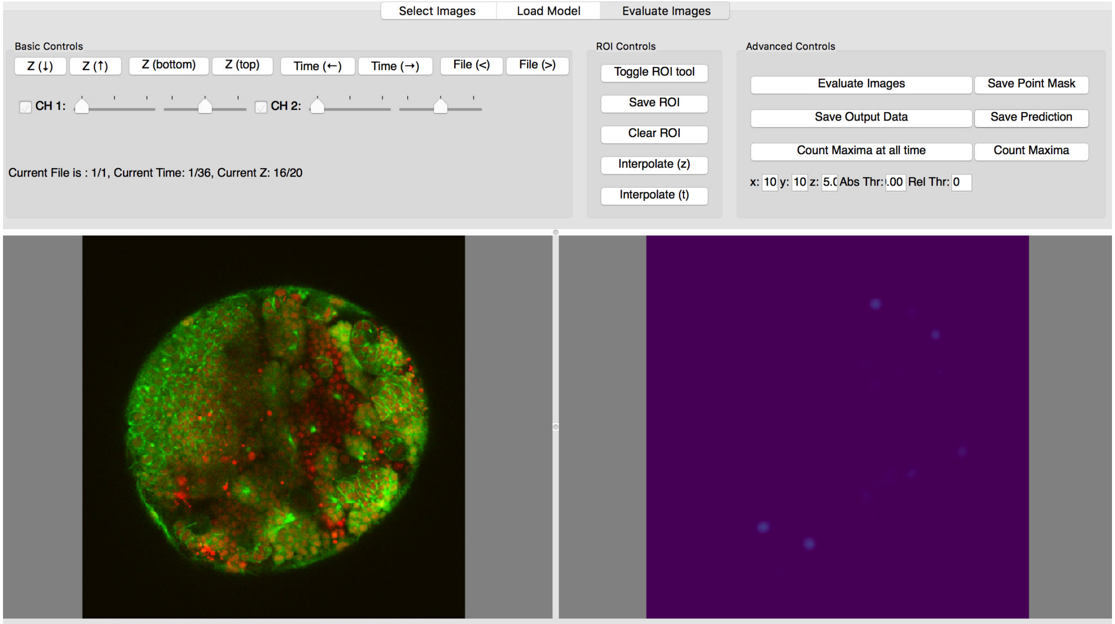


25. When evaluation is complete, you can export the probability maps as tiff files **'Save Prediction'.** Note: this will this be a **hyperstack** if the **input data** is a hyperstack.
26. If you want the counts and cell centre coordinates, use '**Count Maxima at all time**', (this make take a few minutes), and then use **'Save Output Data'** to save the results as excel compatible (.csv) files. You get one file per image file, with the x,y,z coordinate outputs, and one additional file with total counts per image file.
27. If you want to restrict your analysis to a subregion of your image, use the **ROI tool**.

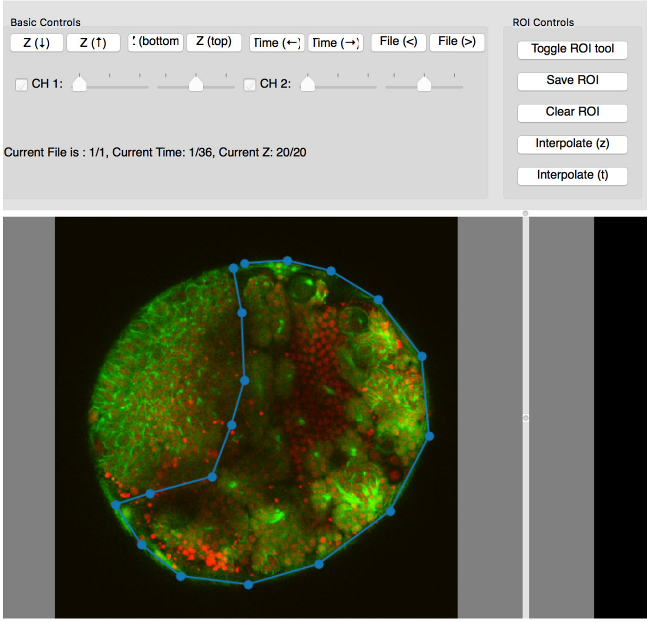

28. First **Toggle ROI tool**
29. Next click a series of points in the current z plane.
30. **Save ROI**
31. Go to another Z plane and click another series of points.

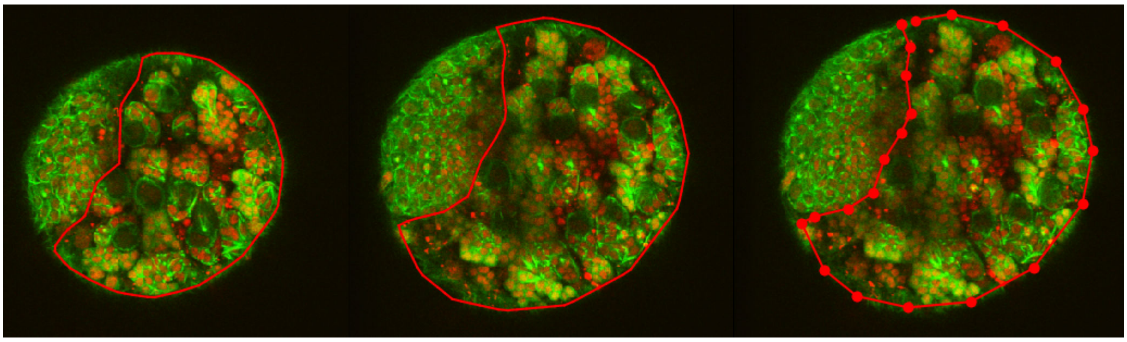

32. Click **Save ROI**
33. Click **Interpolate (z),** to automatically estimate a 3D region of interest between the defined planes
34. Check you're happy with this ROI. If you like you can edit the ROI by adding additional points in more Z planes.
35. If you have images with a time dimension, you can also interpolate between ROIs across T.
36. Repeat for additional files, if necessary.
37. use '**Count Maxima at all time**' to find the counts and centres in these subregions

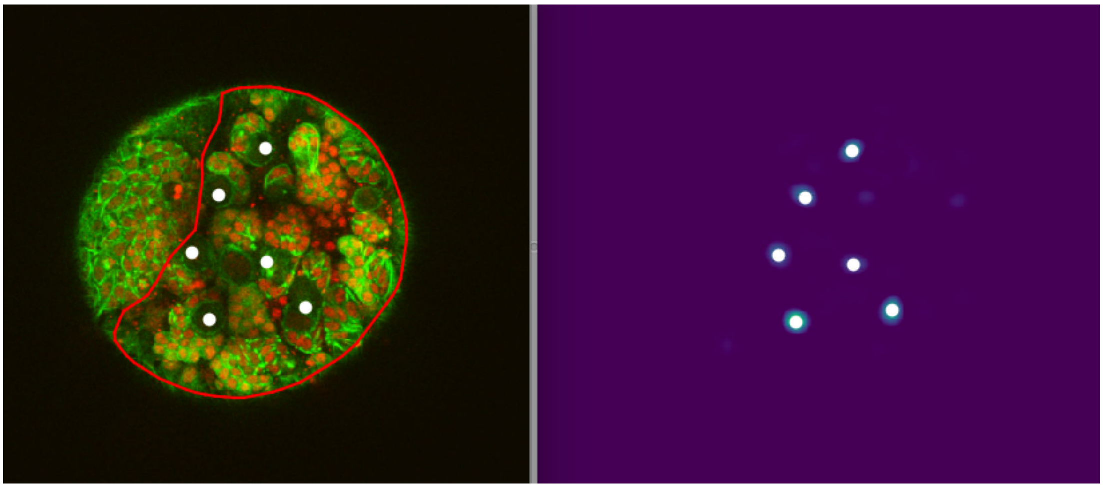

38. Use '**Save Output Data**' to save the results

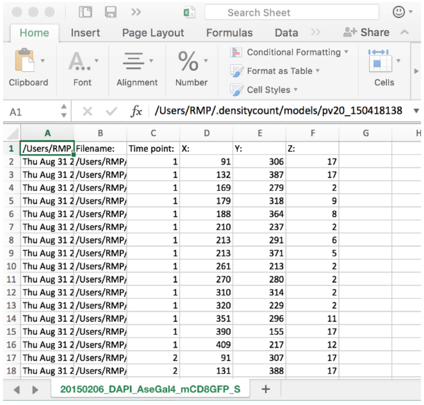


| | A | B | C | D | E | F |
|---|---|---|---|---|---|---|
| 1 | /Users/RMP. | Filename: | Time point: | X: | Y: | Z: |
| 2 | Thu Aug 31 2 | /Users/RMP/ | 1 | 91 | 306 | 17 |
| 3 | Thu Aug 31 2 | /Users/RMP/ | 1 | 132 | 387 | 17 |
| 4 | Thu Aug 31 2 | /Users/RMP/ | 1 | 169 | 279 | 2 |
| 5 | Thu Aug 31 2 | /Users/RMP/ | 1 | 179 | 318 | 9 |
| 6 | Thu Aug 31 2 | /Users/RMP/ | 1 | 188 | 364 | 8 |
| 7 | Thu Aug 31 2 | /Users/RMP/ | 1 | 210 | 237 | 2 |
| 8 | Thu Aug 31 2 | /Users/RMP/ | 1 | 213 | 291 | 6 |
| 9 | Thu Aug 31 2 | /Users/RMP/ | 1 | 213 | 371 | 5 |
| 10 | Thu Aug 31 2 | /Users/RMP/ | 1 | 261 | 213 | 2 |
| 11 | Thu Aug 31 2 | /Users/RMP/ | 1 | 270 | 280 | 2 |
| 12 | Thu Aug 31 2 | /Users/RMP/ | 1 | 310 | 314 | 2 |
| 13 | Thu Aug 31 2 | /Users/RMP/ | 1 | 320 | 229 | 2 |
| 14 | Thu Aug 31 2 | /Users/RMP/ | 1 | 351 | 296 | 11 |
| 15 | Thu Aug 31 2 | /Users/RMP/ | 1 | 390 | 155 | 17 |
| 16 | Thu Aug 31 2 | /Users/RMP/ | 1 | 409 | 217 | 12 |
| 17 | Thu Aug 31 2 | /Users/RMP/ | 2 | 91 | 307 | 17 |
| 18 | Thu Aug 31 2 | /Users/RMP/ | 2 | 131 | 388 | 17 |
